# Matrix stiffness shapes immunometabolic states and response to PD-1 blockade in gastric cancer

**DOI:** 10.64898/2026.10.25.761609

**Authors:** Qiuai Shu, Jian Wu, Ziwei Wang, Yutong Cheng, Xiru Liang, Yingqi Li, Haiqiang Tao, Haitao Shi, Yuxin Peng, Bin Qin, Baicang Zou, Feng Xu, Jinhai Wang, Yongzhan Nie, Yongquan Shi, Daiming Fan, Na Liu, Lu Li, Ting Wen, Ning Xie

## Abstract

The extracellular matrix of solid tumors exhibits spatially heterogeneous stiffness, yet how matrix stiffness shapes antitumor immunity in gastric cancer (GC) remains unclear. Here, we integrated anti-PD-1 clinical cohorts, GC specimens, single-cell and spatial transcriptomic analyses, a syngeneic mouse model, and tunable-stiffness hydrogels to clarify the immunological consequences of matrix stiffness. High matrix stiffness was associated with collagen accumulation and immune exclusion, while higher stiffness-associated transcriptional scores were related to poorer response to PD-1 blockade. Matrix softening markedly reshaped the immune microenvironment and enhanced the efficacy of PD-1 blockade. Mechanistically, stiff matrices promoted F-actin remodeling and YAP nuclear enrichment, driving LDHA-dependent glycolytic reprogramming in GC cells that further impaired the effector function of CD8⁺ T cells. Intriguingly, matrix softening was accompanied by enhanced IFN-γ-responsive PD-L1 expression in GC cells, consistent with adaptive immune resistance that may contribute to the limited antitumor effects of matrix softening alone. These findings suggest that matrix remodeling alleviates spatial and metabolic constraints on antitumor immunity, providing a preclinical rationale for combining matrix softening with PD-1 blockade in GC.

**Graphical abstract:** 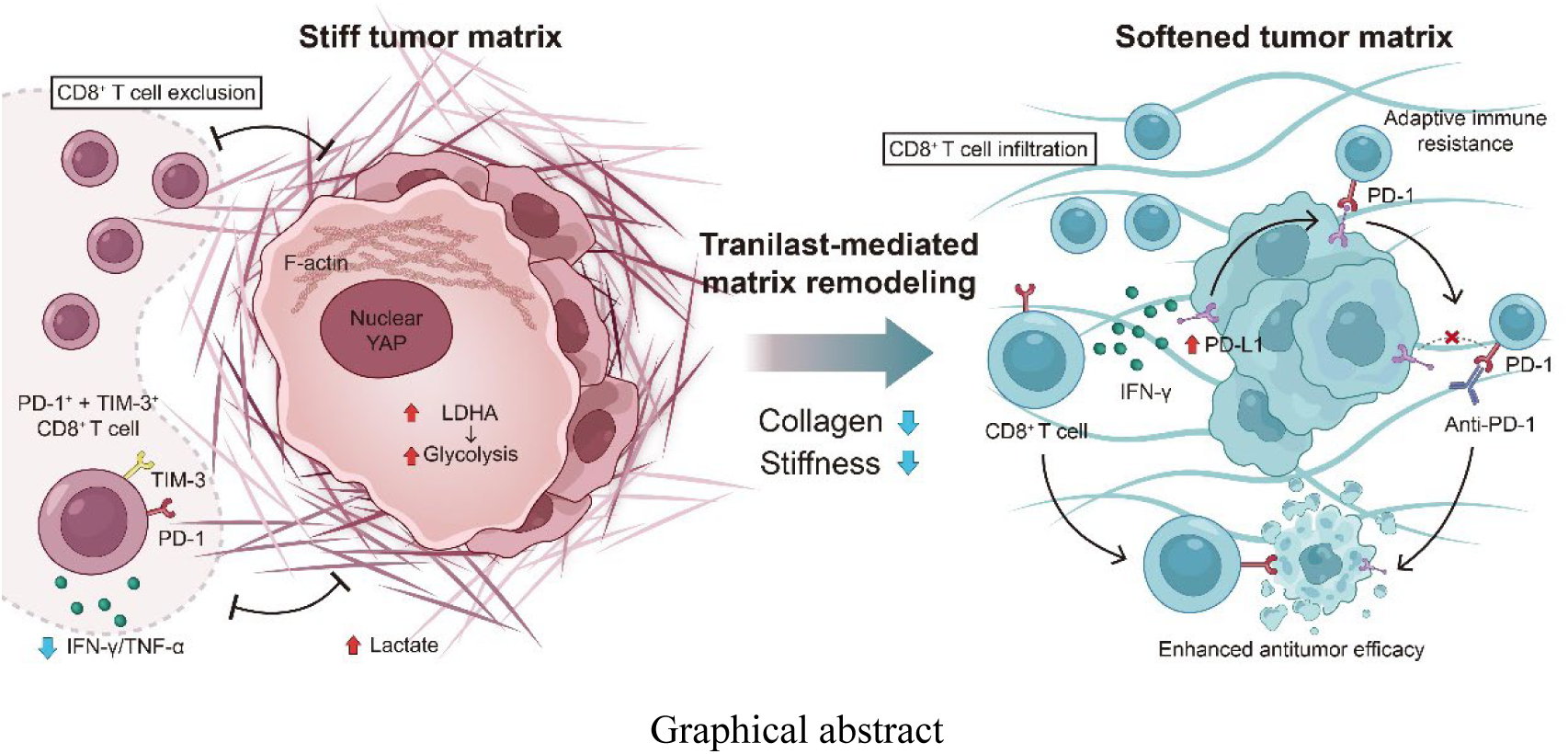

## INTRODUCTION

Immune checkpoint blockade has improved clinical outcomes in advanced gastric cancer (GC), and PD-1 blockade plus chemotherapy has become the standard first-line treatment strategy. Nevertheless, a substantial proportion of patients still fail to achieve durable response (*1, 2*), partly reflecting the pronounced heterogeneity of the tumor immune microenvironment (*3–5*). Beyond the well-studied cellular and biochemical heterogeneity, spatial variation in extracellular matrix (ECM) organization can shape T cell infiltration, providing a physical basis for regional differences in antitumor immunity (*6, 7*). In preclinical models, stromal or matrix normalization can improve immune access and enhance the efficacy of checkpoint blockade (*8–10*). Our previous work characterized the structural and mechanical heterogeneity of the ECM in GC (*11*). However, whether matrix stiffness contributes to immune evasion and limited responses to PD-1 blockade in GC remains unclear.

Beyond its effects on immune cell infiltration, matrix stiffness may also influence antitumor immunity through tumor cell metabolism. Metabolic reprogramming in tumor cells can impair CD8⁺ T cell proliferation and effector function through nutrient deprivation, extracellular acidification, and metabolite accumulation (*12, 13*). Among these metabolic constraints, lactate links tumor glycolysis to T cell dysfunction, as tumor-derived lactate can alter pyruvate utilization and succinate signaling in CD8⁺ T cells, thereby reducing their cytotoxic activity (*14*). Matrix mechanics can regulate glycolysis through actin cytoskeletal organization (*15*), and increased matrix stiffness sustains the Warburg effect to promote pancreatic cancer growth (*16*). In GC, a collagen-rich matrix can increase the production of tumor cell metabolite kynurenic acid, which promotes CAR-T cell exhaustion (*17*), whereas in breast cancer, a stiff fibrotic microenvironment reprograms macrophage arginine metabolism and impairs CD8⁺ T cell function (*18*). Whether matrix stiffness contributes to CD8⁺ T cell dysfunction through altered tumor cell glycolysis in GC remains to be determined.

Here, we show that regional matrix features are associated with immune accessibility and that matrix stiffness modulates the metabolic and immunoregulatory states of GC cells. In clinical cohorts, higher stiffness-associated transcriptional scores were associated with poorer responses to PD-1 blockade, whereas matrix softening increased immune infiltration and markedly enhanced the therapeutic efficacy *in vivo*. Mechanistically, stiff matrices promoted F-actin remodeling and YAP nuclear enrichment, driving LDHA-dependent glycolysis in GC cells and impairing CD8⁺ T cell effector function. By contrast, tumor cells cultured on soft hydrogels showed greater IFN-γ-induced PD-L1 upregulation than those on stiff hydrogels, and matrix softening likewise increased the proportion of PD-L1⁺ tumor cells *in vivo*, suggesting potentially enhanced adaptive immune resistance under low-stiffness conditions. These findings support a model of heterogeneous stiffness-dependent immune evasion and provide a rationale for combining matrix softening with PD-1 blockade to enhance antitumor immunity.

## RESULTS

### Increased matrix stiffness characterizes an immune-excluded tumor microenvironment associated with poor response to PD-1 blockade

To determine whether tumor matrix stiffness is related to clinical response to PD-1 blockade in GC, we analyzed the PRJEB25780 anti-PD-1 cohort (*19*) and integrated the mechanical and immune features in human GC specimens and public single-cell and spatial transcriptomic datasets (Fig. 1A). In the PRJEB25780 anti-PD-1 cohort, stiffness-associated transcriptional scores were significantly higher in non-responders than in responders (Fig. 1B). When patients were stratified by the cohort median, the high-stiffness score group showed a lower response rate (Fig. 1C). Receiver operating characteristic analysis yielded an area under the curve of 0.7146 (fig. S1A). A similar trend was observed in the independent PRJEB40416 cohort, although the difference did not reach statistical significance (*20*) (fig. S1, B and C). Further, we assessed the clinical relevance of stiffness-associated transcriptional score in relation to objective response across both anti-PD-1 cohorts (table S1). Cohort-specific Firth logistic regression yielded odds ratios (ORs) below 1 in both cohorts, indicating a consistent inverse association between stiffness-associated transcriptional score and the odds of objective response (fig. S1D). In the combined individual-patient-data analysis, each 1-SD increase in the within-cohort standardized stiffness-associated transcriptional score was associated with lower odds of objective response to PD-1 blockade (OR, 0.56; 95% confidence interval (CI), 0.29 to 0.97; *P* = 0.038; fig. S1D). The inverse association remained directionally consistent across sensitivity analyses, including covariate adjustment, exclusion of microsatellite instability-high (MSI-H) or Epstein-Barr virus (EBV)-positive cases where applicable, and leave-one-out analyses (fig. S1, E and F; table S1).

**Fig. 1.**
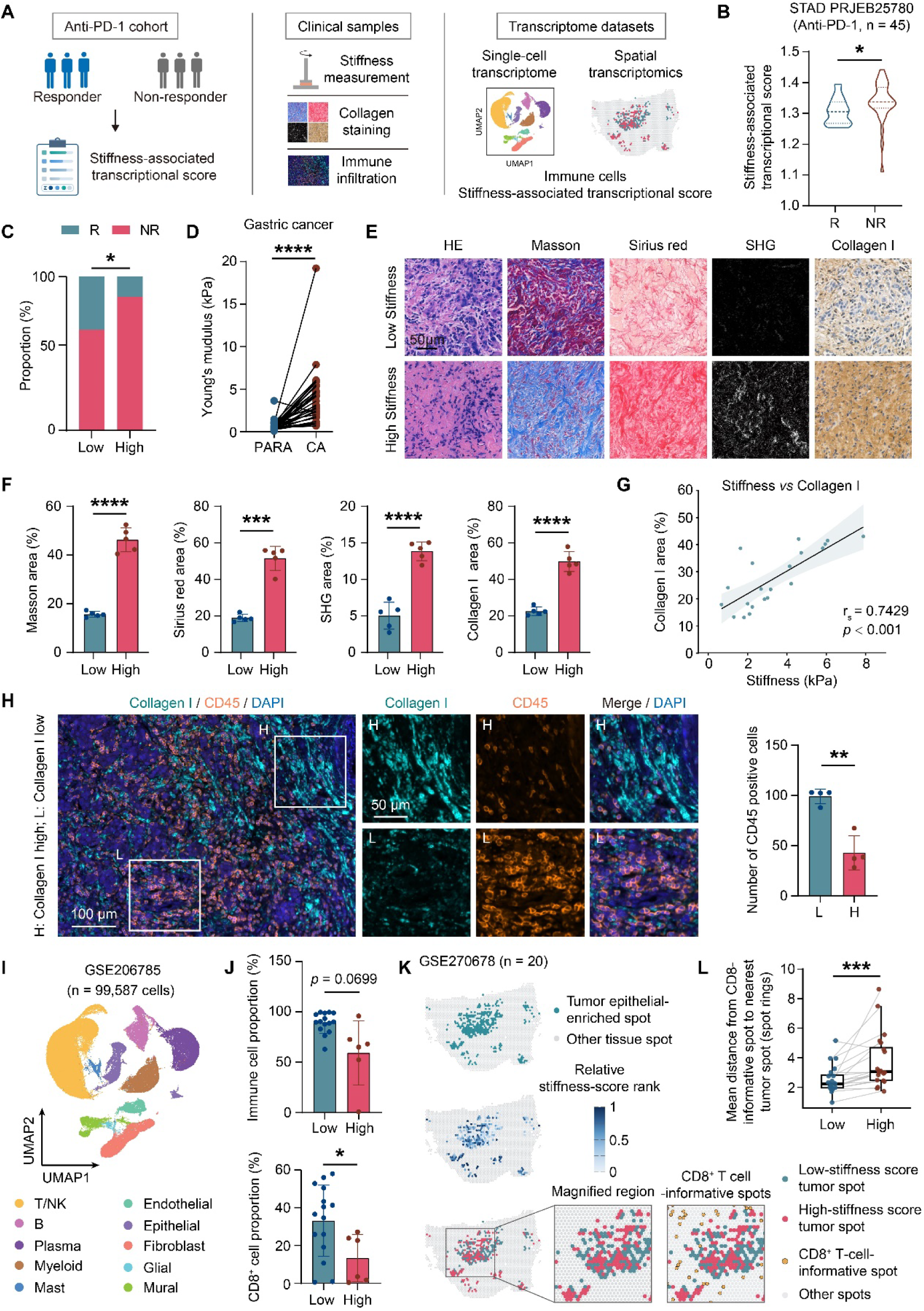
High tumor matrix stiffness is associated with spatial immune exclusion and poor response to PD-1 blockade in gastric cancer. (**A**) Overview of the clinical, tissue, and transcriptomic analyses. (**B**) Stiffness-associated transcriptional scores in responders (R; n = 12) and non-responders (NR; n = 33) in the PRJEB25780 anti-PD-1 cohort. Lines indicate the median and quartiles. (**C**) Response distributions in groups defined by the cohort median stiffness-associated transcriptional score (n = 22 and 23 for low and high groups, respectively). (**D**) Young’s modulus in paired adjacent normal (PARA) and gastric cancer (CA) tissues (n = 34). (**E** and **F**) Representative images of hematoxylin and eosin (H&E), Masson’s trichrome, Sirius red, second harmonic generation (SHG), and collagen I immunohistochemical staining in low- and high-stiffness tumors, with quantification of positive areas (n = 5; points represent patient means across fields). Scale bars, 50 μm. (**G**) Spearman correlation between stiffness and collagen I-positive area (n = 21 patients). (H) Representative multiplex immunofluorescence images of collagen I, CD45, and DAPI in collagen I-low/high regions, with CD45⁺ immune cell quantification by immunohistochemistry on serial sections (n = 4; one point per patient). Scale bars, 100 μm (overview) and 50 μm (magnified). (**I** and **J**) Uniform manifold approximation and projection (UMAP) of the major cell populations in the GSE206785 (n = 99,587 cells) and total immune cell and CD8⁺ T cell proportions in groups defined by the cohort mean stiffness-associated transcriptional score (n = 15 and 6 low- and high-score tumor samples, respectively; one point per sample). (**K**) Representative spatial transcriptomic maps of tumor epithelial-enriched spots, within-slice relative stiffness-score rank, and low/high states defined by the median stiffness-associated transcriptional score among these spots within each slice. The boxed region is magnified with CD8⁺ T cell-informative spots overlaid; other spots are gray. (**L**) Paired patient-level nearest-neighbor distance from CD8⁺ T cell informative spots to low- and high-stiffness score tumor spots, measured in Visium spot rings (n = 20). Distances were averaged within slices, then equally across slices within section groups and across section groups within patients. Each point represents one patient; lines connect pairs. Asterisks indicate unadjusted two-sided Wilcoxon signed-rank *P* values. Benjamini–Hochberg-adjusted *q* values are in table S3. \**P* < 0.05, \*\**P* < 0.01, \*\*\**P* < 0.001, \*\*\*\**P* < 0.0001.

We next characterized the tissue mechanics and ECM organization in human GC specimens. Young’s modulus was markedly higher in GC tissues than in paired adjacent normal tissues (Fig. 1D). High-stiffness tumors also showed greater collagen deposition by Masson’s trichrome staining, Sirius red staining, second harmonic generation (SHG) imaging, and collagen I immunohistochemistry (Fig. 1, E and F), and tissue stiffness positively correlated with collagen I-positive area across specimens (rₛ = 0.7429, *P* < 0.001; Fig. 1G). Given this close association, collagen abundance was used as a histological proxy for relative tissue stiffness in subsequent analyses. Regional analysis within individual tumors showed greater collagen deposition and fewer CD45⁺ immune cells in collagen I-high regions than in collagen I-low regions (Fig. 1H; fig. S1, G and H).

These findings were supported by analysis of the GSE206785 GC single-cell RNA sequencing (scRNA-seq) dataset (*4*). A total of 99,587 cells were annotated into 10 major cell populations, and T/NK cells were subsequently reclustered to identify CD8⁺ T cells (Fig. 1I; fig. S2, A to C). Tumors in the high-stiffness-score group showed a trend toward a lower overall immune cell proportion (*P* = 0.0699) and a significantly reduced CD8⁺ T cell proportion (Fig. 1J). Across samples, stiffness-associated transcriptional scores inversely correlated with both total immune cell proportion (rₛ = −0.5442, *P* = 0.0108) and CD8⁺ T cell proportion (rₛ = −0.5221, *P* = 0.0152; fig. S2, D and E).

To examine these relationships at the spatial level, we analyzed the GSE270678 GC spatial transcriptomic dataset (*21*), which comprised 20 patients and 72 spatial transcriptomic slices (70 evaluable for stiffness score analysis; table S2). Spatial mapping revealed substantial intratumoral heterogeneity in stiffness-associated transcriptional score across tumor epithelial-enriched spots, which were stratified into low- and high-stiffness score states within each slice according to the median stiffness-associated transcriptional score (Fig. 1K). Major epithelial, stromal, and immune cell populations were resolved by spatial deconvolution (Fig. 1K and fig. S2F). At the patient level, CD8⁺ T cell informative spots were located farther from the nearest high-stiffness score tumor spots than from the nearest low-stiffness score tumor spots (Fig. 1L, table S3), and their fractions within two or three spot rings were also lower around high-stiffness score tumor spots (fig. S2G). Together, these clinical, histological, single-cell, and spatial findings link increased matrix stiffness to reduced immune accessibility and poor response to PD-1 blockade in GC.

### Matrix remodeling rewires tumor immune microenvironment and sensitizes tumors to PD-1 blockade

To determine whether tumor matrix softening could improve the efficacy of PD-1 blockade *in vivo*, we used tranilast, an antifibrotic agent reported to reduce collagen deposition and tumor stiffness (*22*), in a subcutaneous GC model in 615 mice and evaluated its effect when combined with anti-PD-1 treatment (Fig. 2A). Interestingly, tranilast alone had little inhibitory effect on tumor growth, whereas anti-PD-1 treatment reduced tumor burden, and the combination markedly suppressed tumor growth and yielded the lowest endpoint tumor weight (Fig. 2, B to D). An additional analysis of log-transformed endpoint relative tumor volume supported a synergistic interaction between tranilast and anti-PD-1 at day 21 (interaction ratio, 0.168; 95% CI, 0.063–0.450; *P* = 0.00145; table S4). The exponential growth rate (eGR)-based Bliss analysis of longitudinal tumor growth also yielded a positive synergy estimate, although the 95% CI marginally crossed zero (synergy score, 41.39; 95% CI, −0.0025 to 144.84; one-sided bootstrap *P* = 0.025; table S4).

**Fig. 2.**
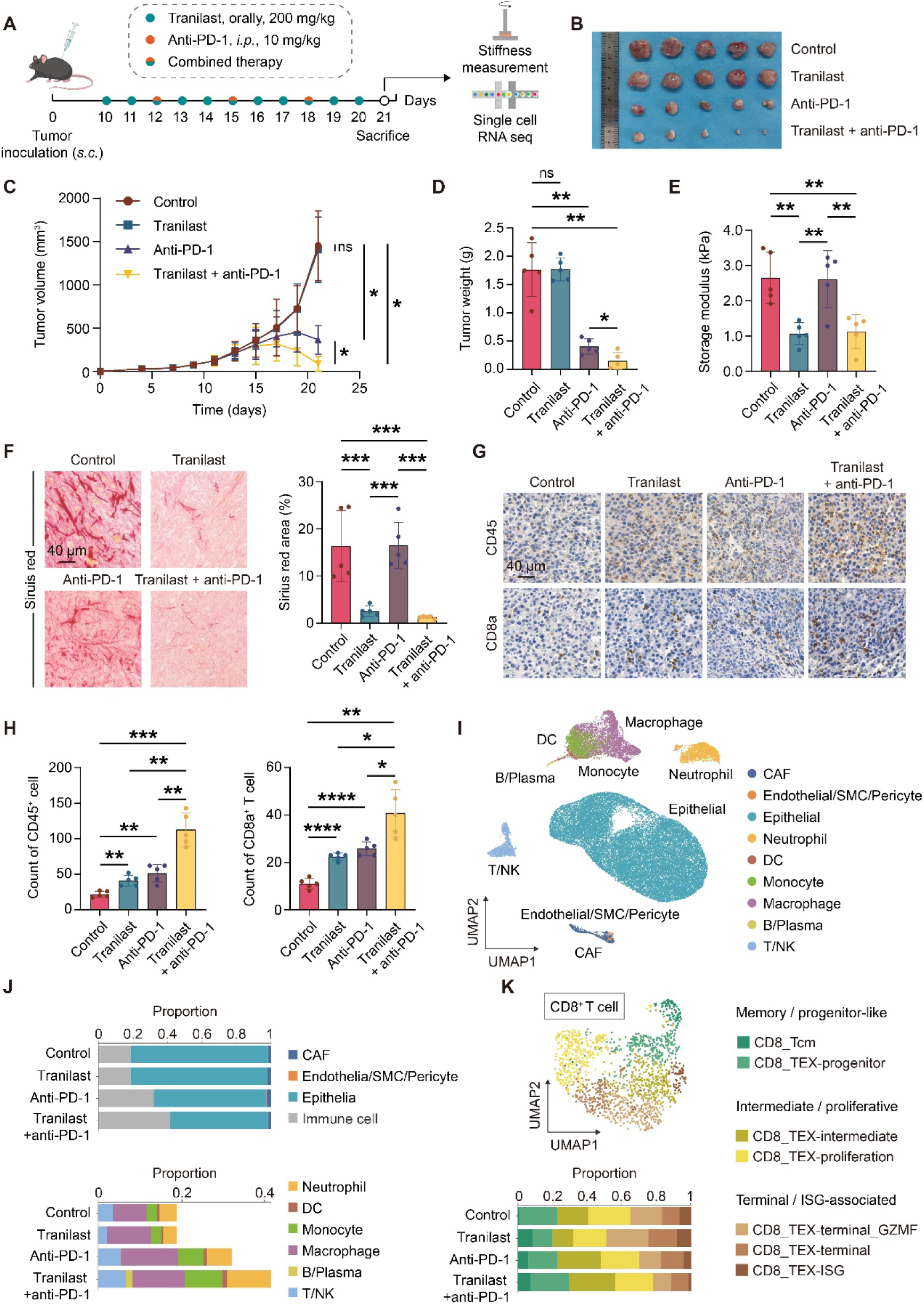
Tumor matrix softening increases immune infiltration and enhances the efficacy of PD-1 blockade in gastric cancer. (**A**) Treatment design in the 615 mouse subcutaneous model: control, tranilast, anti-PD-1, and combination. (**B**) Representative tumor images. (**C**) Tumor growth curves (n = 5). Asterisks indicate comparisons at the final time point. (**D**) Endpoint tumor weights (n = 5; one point per mouse). (**E**) Endpoint tumor storage modulus (n = 5; one point per mouse). (**F**) Sirius red staining and quantification of positive area (n = 5; one point per mouse). Scale bars, 40 μm. (**G** and **H**) Immunohistochemical images of CD45 and CD8α and quantification of CD45⁺ immune cell and CD8⁺ T cell infiltration (n = 5; one point per mouse). Scale bars, 40 μm. (**I** and **J**) UMAP and relative cell population proportions from tumor scRNA-seq (n = 2 mice/group; one sample per mouse). (**K**) Seven CD8⁺ T cell subclusters grouped into memory/progenitor-like, intermediate/proliferative, and terminal/interferon-stimulated gene (ISG)-associated states, with the relative proportions of these states shown. ns, *P* > 0.05, \**P* < 0.05, \*\**P* < 0.01, \*\*\**P* < 0.001, \*\*\*\**P* < 0.0001.

Furthermore, tumor storage modulus was significantly lower in the tranilast and combined treatment groups than in the control and anti-PD-1 monotherapy groups (Fig. 2E), and this softening was accompanied by decreased collagen deposition on Sirius red staining (Fig. 2F), Masson’s trichrome staining, and collagen I immunohistochemistry (fig. S3, A and B). Consistent with the reduced stiffness and collagen deposition, tranilast-induced matrix softening was accompanied by increased infiltration of CD45⁺ immune cells and CD8⁺ T cells, with the highest levels in the combined treatment group (Fig. 2, G and H). No obvious histopathological abnormalities were observed in the major organs across treatment groups (fig. S3C).

We next conducted scRNA-seq to characterize treatment-associated changes in the tumor microenvironment. Major epithelial, stromal, and immune cell populations were identified across the four treatment groups by canonical marker expression (Fig. 2I and fig. S3D). The relative proportions of major cell populations and immune cell subsets varied across treatment groups (Fig. 2J). Reclustering of CD8⁺ T cells resolved seven subclusters, which were grouped into three major states based on transcriptional features: memory/progenitor-like, intermediate/proliferative, and terminal/interferon-stimulated gene (ISG)-associated states (Fig. 2K and fig. S3E). The distribution of CD8⁺ T cell states differed across treatment groups. Specifically, compared with monotherapy, combined treatment increased the proportions of CD8⁺ T central memory (Tcm) cells and CD8⁺ T exhaustion (TEX)-progenitor cells and decreased the proportions of terminal and ISG-associated CD8⁺ TEX cells. Previous studies have reported that memory-like and progenitor-exhausted CD8⁺ T cells are associated with anti-PD-1 response (*23, 24*), whereas terminal CD8⁺ TEX cells and exhausted CD8⁺ T cells with high ISG expression are associated with poor response to anti-PD-1 therapy (*24, 25*). Collectively, these findings indicate that matrix softening reprograms the tumor immune microenvironment and substantially enhances the antitumor efficacy of anti-PD-1 treatment.

### Matrix stiffness reshapes gastric cancer cell metabolism and influences CD8⁺ T cell **function**

Our findings linked collagen accumulation and matrix stiffening to immune exclusion in GC, yet the effects of matrix mechanics on antitumor immunity can extend beyond physical restriction of immune cell access. Previous work has shown that mechanical cues can reprogram malignant epithelial cells, with downstream effects on immune cell behavior and function (*18, 26*). To investigate whether matrix remodeling also altered GC cell programs that could influence CD8^+^ T cell effector function, we therefore focused on the tumor epithelial compartment of the mouse scRNA-seq dataset (Fig. 3A). Gene Ontology (GO) analyses of downregulated genes identified glycolytic process among the top enriched terms in both the tranilast versus control and tranilast plus anti-PD-1 versus anti-PD-1 comparisons (Fig. 3B). Quantitative set analysis for gene expression (QuSAGE) further showed reduced activity of glycolysis/gluconeogenesis and several related metabolic pathways after matrix softening (Fig. 3C). These findings prompted us to further examine the effect of matrix stiffness on tumor glycolytic activity *in vivo* and *in vitro* (Fig. 3D).

**Fig. 3.**
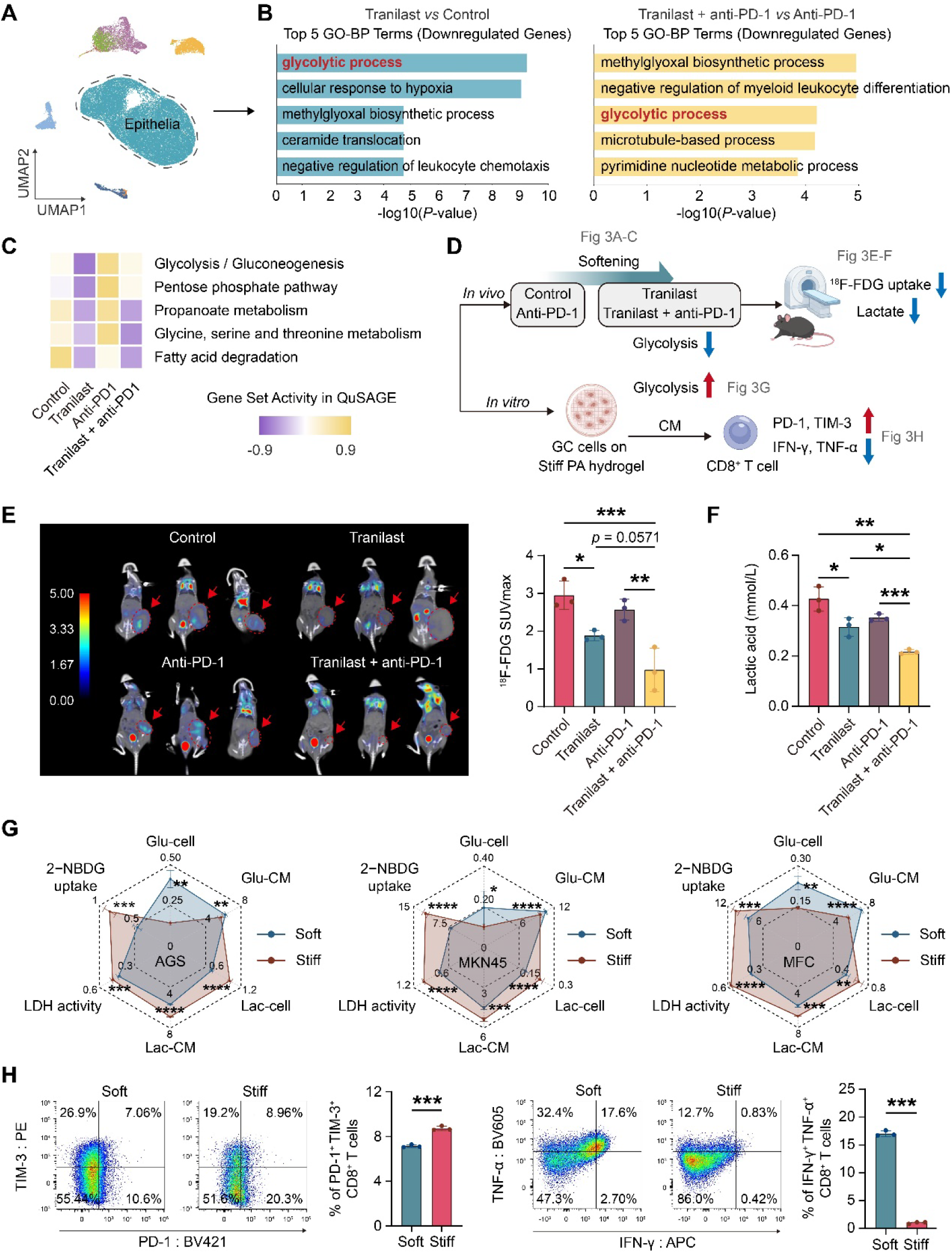
High matrix stiffness promotes glycolytic reprogramming in gastric cancer cells and impairs CD8⁺ T cell effector function. (**A**) UMAP of the scRNA-seq dataset with epithelial cells highlighted. (**B**) Top five Gene Ontology biological process (GO-BP) terms enriched among genes downregulated in epithelial cells in the tranilast versus control and combination versus anti-PD-1 comparisons. (**C**) Metabolic pathway activities across treatment groups assessed by quantitative set analysis for gene expression (QuSAGE). (**D**) Schematic of the experimental strategy used to validate the metabolic changes identified by single-cell analysis. (**E**) ¹⁸F-FDG PET/CT images with maximum standardized uptake value (SUVmax) quantification (n = 3; one point per mouse). (**F**) Quantification of tumor lactate levels (n = 3; one point per mouse). (**G**) Radar plots of glycolytic measurements in AGS, MKN45, and MFC cells on soft or stiff polyacrylamide (PA) hydrogels (n = 4): 2-NBDG uptake, intracellular glucose (Glu-cell), residual medium glucose (Glu-CM), intracellular lactate (Lac-cell), medium lactate (Lac-CM), and cellular LDH activity. Each measurement and cell line has a separate axis scale; units are provided in fig. S4B. (**H**) IFN-γ⁺TNF-α⁺ and PD-1⁺TIM-3⁺ fractions among CD8⁺ T cells exposed to CM from MFC cells cultured on soft or stiff PA hydrogels (n = 3). \**P* < 0.05, \*\**P* < 0.01, \*\*\**P* < 0.001, \*\*\*\**P* < 0.0001.

Consistent with the transcriptomic findings, ¹⁸F-FDG PET/CT showed lower tumor glucose uptake after matrix softening, with the lowest maximum standardized uptake value (SUVmax) in the combined treatment group (Fig. 3E). Tumor lactate levels showed a similar pattern, declining after tranilast and further after combination therapy (Fig. 3F). These *in vivo* findings supported a reduction in tumor glycolytic activity accompanying matrix remodeling. As collagen deposition and tissue stiffness changed in parallel *in vivo*, we complemented these observations using polyacrylamide (PA) hydrogels, in which matrix stiffness could be varied independently of collagen remodeling. AGS, MKN45, and MFC GC cells were cultured on soft or stiff PA hydrogels. Rotational rheometry confirmed the stiffness difference between the two PA hydrogel substrates (fig. S4A). Cells cultured on stiff PA hydrogels consistently showed increased 2-NBDG uptake and lower residual glucose in the conditioned medium (CM), together with increased intracellular and extracellular lactate and higher lactate dehydrogenase (LDH) activity (Fig. 3G and fig. S4B). Bulk RNA sequencing (bulk RNA-seq) further showed enrichment of canonical glycolysis programs in cells on stiff PA hydrogels (fig. S4C). Consistent metabolic changes across three cell lines supported a direct effect of matrix stiffness on the glycolytic phenotype of GC cells.

We then tested whether matrix stiffness altered the capacity of GC cells to modulate CD8⁺ T cell function. Mouse splenic lymphocytes were cultured for 24 h with CM from GC cells cultured on soft or stiff PA hydrogels. Compared with CM from cells on soft hydrogels, CM from cells on stiff PA hydrogels increased the proportion of PD-1⁺TIM-3⁺ CD8⁺ T cells and markedly reduced the proportion of IFN-γ⁺TNF-α⁺ CD8⁺ T cells (Fig. 3H; gating strategy in fig. S5). These findings support an indirect suppressive effect of matrix stiffness on CD8⁺ T cell effector function through GC cell-derived CM; whether the accompanying metabolic reprogramming contributed to this effect was examined further.

### LDHA mediates the glycolytic response to high matrix stiffness and contributes to suppression of CD8⁺ T cell function

To identify candidate metabolic mediators of stiffness-associated CD8⁺ T cell suppression, we integrated differential expression results from the two *in vivo* tumor epithelial cell comparisons with bulk RNA-seq data from AGS cells cultured on soft or stiff PA hydrogels. LDHA and AHNAK were the only two differentially expressed genes (DEGs) shared across all three comparisons (Fig. 4A). Notably, LDHA was downregulated in both *in vivo* comparisons involving tranilast treatment (fig. S6, A and B). Given this consistent expression pattern and the glycolytic phenotype identified in Fig. 3, we prioritized LDHA for further investigation.

**Fig. 4.**
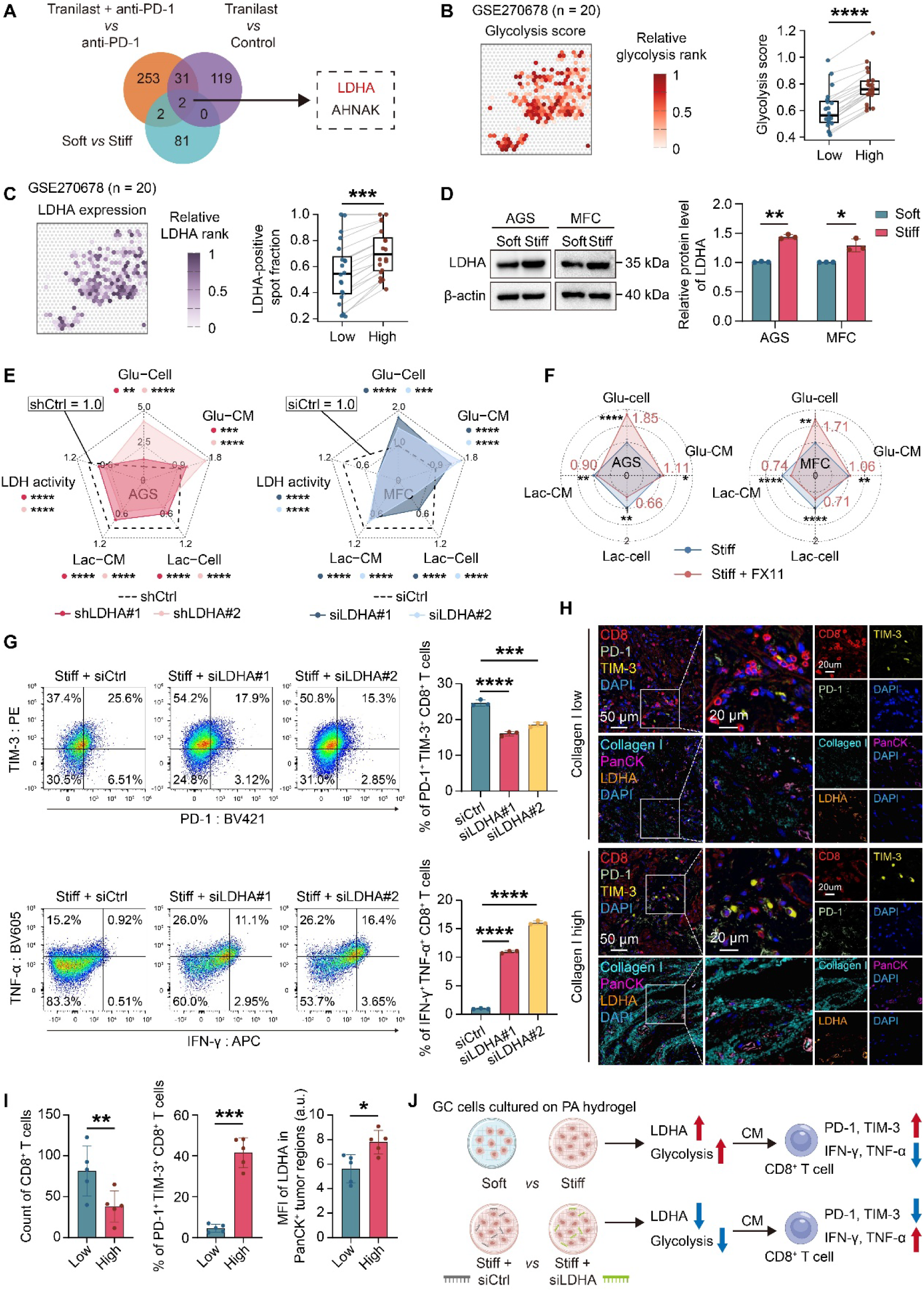
LDHA mediates the glycolytic reprogramming induced by high matrix stiffness and contributes to CD8⁺ T cell functional suppression. (**A**) Overlap of differentially expressed genes (DEGs) from scRNA-seq and bulk transcriptomic datasets, identifying LDHA and AHNAK. (**B** and **C**) Spatial maps of glycolysis scores (**B**) and LDHA expression and positivity (**C**), with patient-level paired comparisons of glycolysis scores and LDHA-positive spot fractions between low- and high-stiffness-score tumor spots defined in Fig. 1. Asterisks denote unadjusted two-sided Wilcoxon signed-rank *P* values; Benjamini–Hochberg-adjusted *q* values are in table S3. (**D**) Representative LDHA Western blots and quantification in AGS and MFC cells on soft or stiff PA hydrogels (n = 3). (**E**) Radar plots of glycolytic measurements in AGS and MFC cells cultured on stiff PA hydrogels after LDHA knockdown (n = 3): intracellular glucose (Glu-cell), residual medium glucose (Glu-CM), intracellular lactate (Lac-cell), medium lactate (Lac-CM), and cellular LDH activity. Each measurement was normalized to the corresponding shCtrl or siCtrl group, set to 1.0. (**F**) Radar plots of glycolytic measurements in AGS and MFC cells cultured on stiff PA hydrogels after pharmacological inhibition of LDHA with FX11 (n = 4): Glu-cell, Glu-CM, Lac-cell, and Lac-CM, as defined in (**E**). Each measurement was normalized to the corresponding vehicle-treated control on stiff PA hydrogels, set to 1.0. (**G**) IFN-γ⁺TNF-α⁺ and PD-1⁺TIM-3⁺ fractions among CD8⁺ T cells exposed to CM from control or LDHA-knockdown MFC cells on stiff PA hydrogels (n = 3). (**H**) Representative multiplex immunofluorescence of collagen I, pan-cytokeratin (PanCK), LDHA, CD8, PD-1, TIM-3, and DAPI in low/high collagen I regions of human GC specimens. Scale bars, 50 μm (overview) and 20 μm (magnified). (**I**) Quantification of CD8⁺ T cell numbers, PD-1⁺TIM-3⁺ fractions among CD8⁺ T cells, and LDHA mean fluorescence intensity in PanCK⁺ tumor regions shown in (**H**) (n = 5; one point per patient). (**J**) Schematic linking matrix stiffness, LDHA-dependent glycolysis, and CD8⁺ T cell suppression. \**P* < 0.05, \*\**P* < 0.01, \*\*\**P* < 0.001, \*\*\*\**P* < 0.0001.

Spatial transcriptomic analysis provided additional support for this association in human GC. Among the tumor spots classified by stiffness-associated transcriptional scores in Fig. 1, high-stiffness score tumor spots showed higher glycolysis scores and a greater fraction of LDHA-positive spots than low-stiffness score tumor spots. However, LDHA expression among LDHA-positive tumor spots was lower in the high-stiffness score group and was negatively correlated with stiffness-associated transcriptional score in the continuous analysis (Fig. 4, B and C; table S3). Public transcriptomic datasets also showed higher LDHA mRNA expression in GC than in normal gastric tissues, and paired immunohistochemistry confirmed increased LDHA protein expression in tumor tissues compared with adjacent normal tissues (fig. S6, C and D). In parallel, LDHA protein expression was higher in AGS and MFC cells cultured on stiff PA hydrogels than in cells on soft hydrogels (Fig. 4D) with similar increases in MKN45 cells by western blotting and across all three cell lines by immunofluorescence (fig. S6, E and F).

The functional role of LDHA in the glycolytic response to high matrix stiffness was examined using genetic and pharmacological approaches. LDHA knockdown in AGS, MKN45, and MFC cells on stiff PA hydrogels increased residual glucose levels in the CM and reduced intracellular and extracellular lactate levels and LDH activity (Fig. 4E and fig. S7). Pharmacological inhibition of LDHA with FX11 similarly increased residual glucose levels in the CM and reduced intracellular and extracellular lactate levels across all three cell lines (Fig. 4F and fig. S8). These findings support a functional contribution of LDHA to the enhanced glycolytic phenotype of GC cells induced by high matrix stiffness.

We then examined whether tumor cell LDHA contributed to the suppressive effect of CM from GC cells cultured on stiff PA hydrogels. Mouse splenic lymphocytes were exposed to CM for 24 h from MFC cells cultured on stiff PA hydrogels with or without LDHA knockdown. LDHA depletion reduced the proportion of PD-1⁺TIM-3⁺ CD8⁺ T cells and increased the proportion of IFN-γ⁺TNF-α⁺ CD8⁺ T cells compared with control CM (Fig. 4G). Thus, disrupting LDHA in GC cells not only attenuated the glycolytic response to high matrix stiffness but also alleviated the suppressive effect of their CM on CD8⁺ T cell effector function.

The spatial associations among matrix composition, tumor cell LDHA expression, and CD8⁺ T cell phenotypes were further evaluated in human GC tissues. Multiplex immunofluorescence showed fewer CD8⁺ T cells and a higher proportion of PD-1⁺TIM-3⁺ cells among CD8⁺ T cells in regions with high collagen I abundance (Fig. 4, H and I), where LDHA expression in pan-cytokeratin (PanCK)⁺ tumor cells was also higher (Fig. 4, H and I). Together, these findings identify LDHA as an important mediator of the stiffness-induced glycolytic response in GC cells and support its contribution to the suppression of CD8⁺ T cell effector function (Fig. 4J).

### Matrix stiffness regulates LDHA through F-actin remodeling and YAP activation

Having established the role of LDHA in stiffness-induced glycolytic reprogramming, we examined how its expression was regulated. Bulk RNA-seq of AGS cells cultured on stiff versus soft PA hydrogels showed increased expression of LDHA and several genes associated with YAP/TAZ transcriptional activity, including ANKRD1, AMOTL2, and CYR61 (fig. S6B). Gene set enrichment analysis (GSEA) further showed enrichment of the Cordenonsi YAP conserved signature and the Kanai YAP/TAZ up signature under stiff conditions (Fig. 5A and fig. S9A). Accordingly, the ratio of Ser127-phosphorylated YAP to total YAP (p-YAP/YAP) was reduced in AGS and MFC cells cultured on stiff PA hydrogels (Fig. 5B), and immunofluorescence showed increased nuclear YAP enrichment (Fig. 5, C and D). Similar changes were observed in MKN45 cells (fig. S9, B and C).

**Fig. 5.**
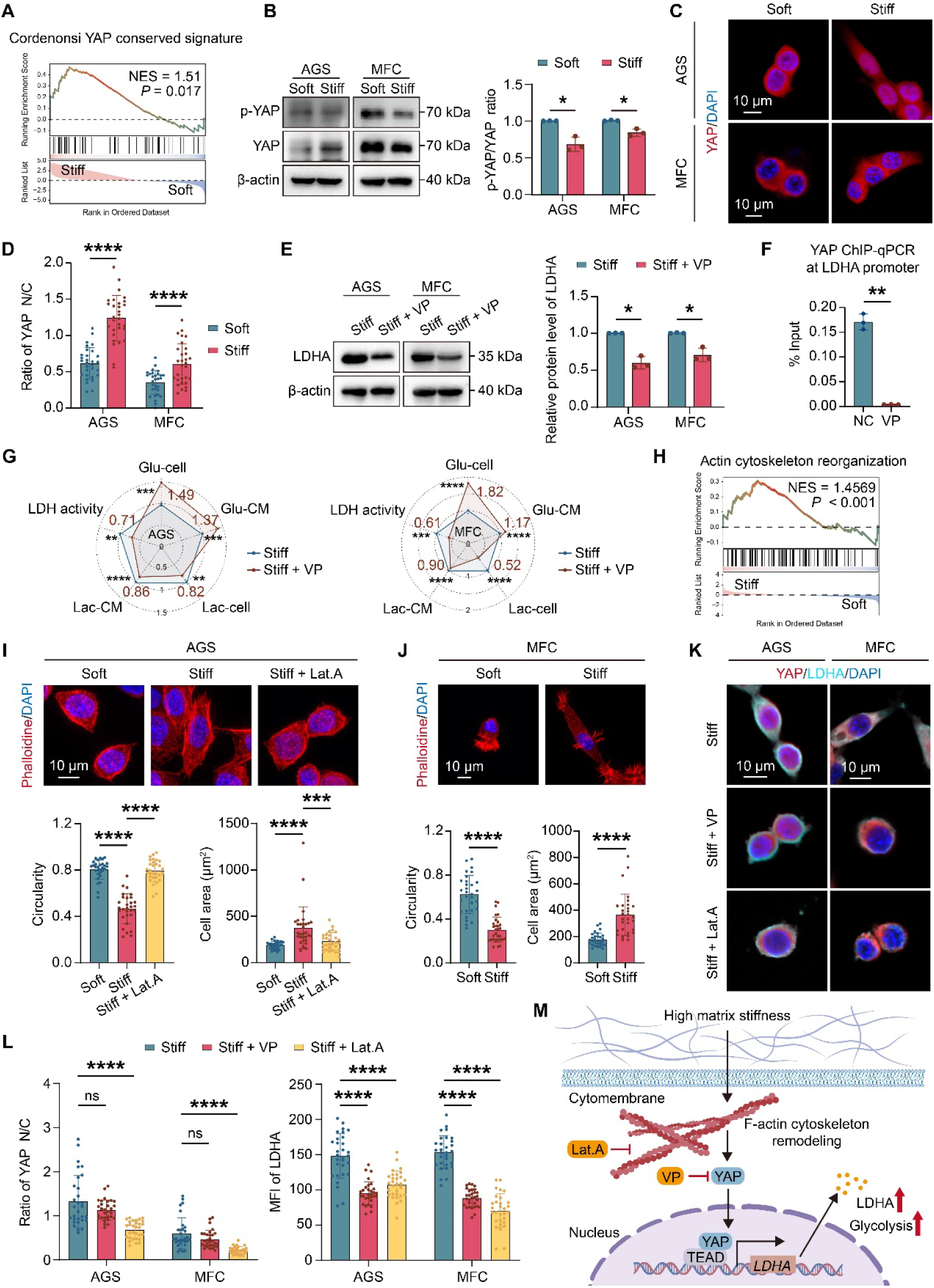
High matrix stiffness activates YAP through F-actin remodeling to promote LDHA expression and glycolytic reprogramming. (**A**) Gene set enrichment analysis (GSEA) of bulk RNA-seq data comparing AGS cells on stiff versus soft PA hydrogels for the Cordenonsi YAP conserved signature. (**B**) Representative Western blots of Ser127-phosphorylated YAP (p-YAP), total YAP, and β-actin, with p-YAP/YAP quantification in AGS and MFC cells on soft or stiff PA hydrogels (n = 3). (**C** and **D**) Representative YAP immunofluorescence (**C**) and nuclear-to-cytoplasmic intensity ratio (**D**) in AGS and MFC cells on soft or stiff hydrogels; nuclei were counterstained with DAPI. (N = 3 independent experiments, n = 30 cells per condition). Scale bars, 10 μm. (**E**) Representative LDHA Western blots and quantification in AGS and MFC cells on stiff PA hydrogels with vehicle or verteporfin (VP) (n = 3). (**F**) YAP chromatin immunoprecipitation followed by quantitative PCR (ChIP-qPCR) at the LDHA promoter in AGS cells treated with vehicle or VP (n = 3), expressed as the percentage of input DNA. (**G**) Radar plots of glycolytic measurements in AGS and MFC cells cultured on stiff PA hydrogels with vehicle or VP (n = 4): intracellular glucose (Glu-cell), residual medium glucose (Glu-CM), intracellular lactate (Lac-cell), medium lactate (Lac-CM), and cellular LDH activity. Each measurement was normalized to the corresponding vehicle control, set to 1.0. (**H**) GSEA of bulk RNA-seq data comparing AGS cells on stiff versus soft PA hydrogels for actin cytoskeleton reorganization. (**I** and **J**) Representative phalloidin staining and quantification of cell area and circularity in AGS cells on soft or stiff hydrogels or stiff hydrogels with latrunculin A (Lat.A) (**I**), and MFC cells on soft or stiff hydrogels (**J**). (N = 3 independent experiments, n = 30 cells per condition). Scale bars, 10 μm. (**K** and **L**) Representative YAP and LDHA immunofluorescence (**K**), with YAP nuclear-to-cytoplasmic ratios and LDHA mean fluorescence intensity (**L**), in AGS and MFC cells on stiff hydrogels with vehicle, VP, or Lat.A. (N = 3 independent experiments, n = 30 cells per condition). Scale bars, 10 μm. (**M**) Schematic of the F-actin–YAP–LDHA pathway. TEAD is shown as YAP’s canonical transcriptional partner; its role in LDHA regulation was not directly tested. ns, *P* > 0.05, \**P* < 0.05, \*\**P* < 0.01, \*\*\**P* < 0.001, \*\*\*\**P* < 0.0001.

We next examined whether YAP contributed to LDHA upregulation under stiff conditions. YAP silencing reduced LDHA protein expression in AGS and MKN45 cells cultured on stiff PA hydrogels (fig. S9D), and verteporfin (VP) treatment similarly decreased LDHA expression in AGS, MKN45, and MFC cells (Fig. 5E and fig. S9E). In addition, YAP1 and LDHA mRNA expression were positively correlated in public GC transcriptomic data (fig. S9F). Chromatin immunoprecipitation followed by quantitative PCR (ChIP-qPCR) further showed YAP enrichment at the LDHA promoter that was markedly reduced by VP treatment (Fig. 5F). These results supported YAP-dependent transcriptional regulation of LDHA. Consistent with this, VP increased residual glucose levels in the CM and reduced intracellular and extracellular lactate levels and LDH activity in AGS, MKN45, and MFC cells cultured on stiff PA hydrogels (Fig. 5G and fig. S10A).

Given the role of actin cytoskeletal tension in YAP regulation (*27*), we next examined F-actin remodeling under stiff conditions. GSEA of our bulk RNA-seq data showed enrichment of the actin cytoskeleton reorganization in GC cells cultured on stiff PA hydrogels (Fig. 5H). Phalloidin staining revealed pronounced F-actin reorganization accompanied by increased cell area and reduced circularity in AGS and MFC cells on stiff versus soft PA hydrogels (Fig. 5, I and J). Similar morphological changes were observed in MKN45 cells (fig. S10B). Treatment with latrunculin A (Lat.A) largely reversed these morphological changes in AGS cells (Fig. 5I).

Disruption of F-actin with Lat.A markedly reduced YAP nuclear enrichment and LDHA expression in AGS and MFC cells, whereas VP decreased LDHA expression without significantly altering YAP nuclear localization (Fig. 5, K and L). Western blot analysis further showed that Lat.A increased the p-YAP/YAP ratio and reduced LDHA protein expression in AGS, MKN45, and MFC cells (fig. S9, G and H). Similar effects on YAP nuclear enrichment and LDHA expression were confirmed in MKN45 cells by immunofluorescence (fig. S9I). Together, these findings support a pathway in which stiffness-induced F-actin remodeling promotes YAP activation and subsequent LDHA expression, contributing to glycolytic reprogramming (Fig. 5M).

### Matrix softening modulates IFN-γ-responsive PD-L1 expression in gastric cancer cells

Despite reduced tumor glycolytic activity and increased immune infiltration after matrix softening, tranilast alone exhibited limited tumor control, whereas its combination with PD-1 blockade produced a marked antitumor effect (Fig. 2). Recent evidence indicates that matrix stiffness can also modulate immunoregulatory programs in tumor cells, including PD-L1 expression (*28*). We therefore examined whether matrix remodeling was accompanied by additional immunoregulatory responses in GC cells that might constrain antitumor immunity.

Reclustering of tumor epithelial cells identified six subclusters, one of which showed high *Cd274* expression and was designated the PD-L1-high epithelial state (Fig. 6A; fig. S11, A and B). The PD-L1-high epithelial state was more abundant in anti-PD-1-treated tumors, reached its highest proportion in the combination group, and increased modestly with tranilast alone (Fig. 6B and fig. S11C). Transcriptional profiling further revealed that this cell state showed higher expression of genes involved in type II interferon response, together with genes associated with antigen processing and adaptive immune resistance (Fig. 6C). The coordinated expression of *Stat1, Irf1, Socs1, Cxcl9, Cxcl10,* and *Cd274* indicated an enhanced IFN-γ-responsive program and was consistent with an adaptive immune resistance phenotype linked to local IFN-γ signaling and tumor cell PD-L1 expression (*29, 30*).

**Fig. 6.**
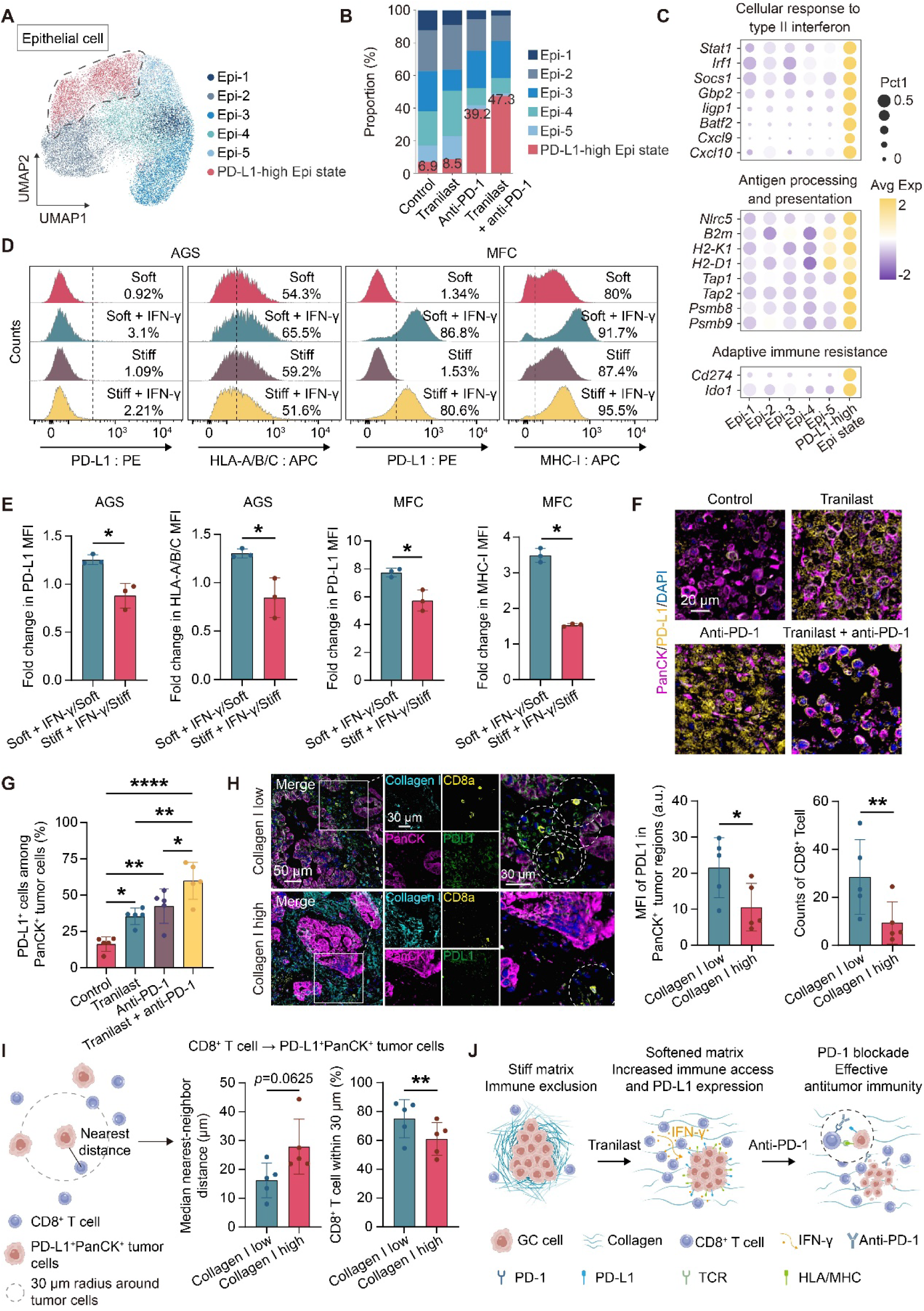
Low matrix stiffness enhances IFN-γ-responsive PD-L1 expression in gastric cancer. (**A** and **B**) UMAP of six tumor epithelial subclusters from scRNA-seq of mouse gastric tumors (Epi-1–Epi-5 and the PD-L1-high epithelial state) and their relative proportions across treatment groups. Proportions were calculated among all epithelial cells in each group; labels indicate the PD-L1-high fraction. (**C**) Dot plots of genes associated with type II interferon response, adaptive immune resistance, and antigen processing and presentation. Dot size indicates the expressing cell fraction; color indicates average expression. (**D** and **E**) Surface PD-L1 and HLA-A/B/C or MHC-I expression by flow cytometry (**D**) and IFN-γ-induced fold changes in median fluorescence intensity (MFI) (**E**) in GC cells on soft or stiff PA hydrogels after 24 h stimulation (n = 3). Flow labels indicate positive cell percentages; fold changes were calculated as stimulated/unstimulated MFI. (**F** and **G**) Representative PanCK and PD-L1 multiplex immunofluorescence (**F**) and percentages of PD-L1⁺ cells among PanCK⁺ tumor cells (**G**) in 615 mouse subcutaneous tumors (n = 5; one point per mouse). Scale bars, 20 μm. (**H**) Representative multiplex immunofluorescence of collagen I, PanCK, PD-L1, and CD8α in low/high collagen I regions of human GC, with quantification of PD-L1 mean fluorescence intensity (MFI) in PanCK⁺ tumor regions and CD8⁺ T cell abundance (n = 5; one point per patient). Scale bars, 50 μm and 30 μm, as indicated. (**I**) Nearest-neighbor analysis of CD8⁺ T cell proximity to PD-L1⁺PanCK⁺ tumor cells, showing median distances and the fraction of CD8⁺ T cells within 30 μm of these tumor cells in low/high collagen I regions (n = 5; one point per patient). (**J**) Proposed model of tranilast-mediated matrix softening increasing immune infiltration and enhancing PD-1 blockade efficacy. Local IFN-γ signaling and adaptive PD-L1 expression may limit the efficacy of softening alone. \**P* < 0.05, \*\**P* < 0.01, \*\*\*\**P* < 0.0001.

To determine whether matrix stiffness modulated this IFN-γ-responsive phenotype, AGS and MFC cells were cultured on soft or stiff PA hydrogels and stimulated with IFN-γ for 24 h. Basal surface PD-L1 positivity was low in both cell lines (Fig. 6D and fig. S11D). After IFN-γ stimulation, the fold change in PD-L1 median fluorescence intensity (MFI) relative to the corresponding unstimulated control was greater on soft than on stiff hydrogels in both AGS and MFC cells (Fig. 6E and fig. S11E). The IFN-γ-induced change in PD-L1 MFI (ΔMFI) was also greater on soft hydrogels (fig. S11F). MFC cells showed a greater IFN-γ-induced fold increase in MHC-I MFI on soft hydrogels. In AGS cells, the fold change in HLA-A/B/C MFI was higher on soft hydrogels but decreased on stiff hydrogels following IFN-γ stimulation, although the difference in ΔMFI did not reach statistical significance (Fig. 6, D and E; fig. S11, D to F). *In vivo*, the proportion of PD-L1⁺ cells among PanCK⁺ tumor cells was higher in both the tranilast and anti-PD-1 groups than in the control group. Combined treatment further increased this proportion compared with either monotherapy (Fig. 6, F and G).

We next examined the spatial relationship among collagen abundance, tumor cell PD-L1 expression, and CD8⁺ T cell distribution in human GC tissues. Using collagen abundance as the histological proxy for matrix stiffness established above, regions with low collagen I abundance showed higher PD-L1 expression in PanCK⁺ tumor cells and contained more CD8⁺ T cells than regions with high collagen I abundance (Fig. 6H). Spatial analysis further showed a trend toward a shorter distance between CD8⁺ T cells and PD-L1⁺PanCK⁺ tumor cells in collagen I-low regions (*P* = 0.0625), and a significantly greater proportion of CD8⁺ T cells located within 30 μm of PD-L1⁺ tumor cells (Fig. 6I). Together, these findings show that low matrix stiffness enhances IFN-γ-responsive PD-L1 expression *in vitro* and that matrix softening is accompanied by increased tumor cell PD-L1 expression *in vivo*. The accompanying PD-L1 response suggests potential adaptive immune resistance, providing a rationale for combining matrix softening with PD-1 blockade (Fig. 6J).

## DISCUSSION

In this study, we found that matrix stiffness was associated with immune accessibility and modulated the metabolic and immunoregulatory states of GC cells. Higher stiffness-associated transcriptional scores were associated with poorer responses to PD-1 blockade. Mechanistically, F-actin remodeling and YAP nuclear enrichment contributed to stiffness-induced increases in LDHA expression and glycolytic activity, and LDHA knockdown reduced the suppressive effect of tumor cell CM on CD8⁺ T cell function. Beyond metabolic regulation, matrix stiffness also modulated the IFN-γ-induced PD-L1 response of GC cells, with greater induction under low-stiffness conditions *in vitro*. *In vivo*, matrix softening improved immune accessibility, reduced tumor glycolytic activity, and was accompanied by increased tumor cell PD-L1 expression. Although matrix softening alone provided limited tumor control, it enhanced the antitumor efficacy of PD-1 blockade. Collectively, these findings support a model in which matrix remodeling alleviates spatial and metabolic constraints on antitumor immunity, while the accompanying PD-L1 response may contribute to adaptive immune resistance.

Effective antitumor immunity depends not only on the presence of T cells within a tumor, but also on their ability to reach malignant cells (*31–34*). Fibrosis and matrix stiffening are increasingly recognized as physical features associated with reduced immune infiltration, whereas experimental matrix softening can increase lymphocyte entry and improve the efficacy of immune checkpoint blockade (*18, 35*). Regional variation in immune infiltration may partly reflect spatial heterogeneity in matrix architecture (*7, 36*). Our findings extend these observations to GC by linking matrix heterogeneity to regional immune accessibility. Collagen abundance was closely associated with measured tissue stiffness, and collagen-rich regions within individual tumors contained fewer immune cells. Spatial transcriptomic analysis further showed that CD8⁺ T cell informative spots were located farther from tumor epithelial-enriched spots with high stiffness-associated transcriptional scores than from those with low scores. Together, these findings support an association between regional matrix differences and the coexistence of immune-infiltrated and immune-restricted regions within individual GC lesions. Consistent with this interpretation, tranilast treatment *in vivo* reduced collagen deposition and tumor stiffness while increasing immune cell infiltration.

Beyond restricting immune access, matrix stiffness can also reprogram tumor cell metabolism (*15, 37*), which in turn shapes local immunity through nutrient consumption and metabolite production that affect T cell function (*13, 38*). In our study, high matrix stiffness induced F-actin remodeling and YAP nuclear enrichment, together with increased LDHA expression and glycolytic activity. Disruption of F-actin or inhibition of YAP reduced LDHA expression, supporting a model in which F-actin remodeling and YAP activation mediate the metabolic response to matrix stiffness. Importantly, the effects extended beyond the tumor cells themselves. CM from MFC cells cultured on stiff matrices impaired CD8⁺ T cell effector function, whereas LDHA knockdown attenuated both the glycolytic phenotype and the suppressive effect of the CM. Consistent with this finding, recent work has shown that LDHA-dependent glycolysis in tumor cells can shape an immunosuppressive metabolic environment and influence immunotherapy response (*39*). Together, these findings extend the immunoregulatory effects of matrix stiffness beyond spatial immune exclusion, demonstrating that stiffness-induced glycolytic reprogramming in tumor cells can also impair CD8⁺ T cell function.

IFN-γ is a central mediator of antitumor T cell responses, promoting antigen presentation and tumor cell recognition, but it can also induce PD-L1 expression and thereby contribute to adaptive immune resistance (*29, 30, 40*). In our scRNA-seq analysis, the PD-L1-high epithelial state expressed IFN-γ-response and antigen processing and presentation genes alongside Cd274, placing PD-L1 expression within a broader IFN-γ-associated transcriptional program. Under the same IFN-γ stimulation, GC cells cultured on soft matrices showed greater changes in surface PD-L1 expression than those on stiff matrices, as assessed by both MFI fold change and ΔMFI. *In vivo*, tranilast treatment increased the proportion of PD-L1-positive tumor cells, with the highest proportion observed after combined treatment with anti-PD-1. Consistent with this pattern, regions with low collagen abundance in human GC contained more CD8⁺ T cells and showed higher PD-L1 expression in tumor cells than regions with high collagen abundance. These observations suggest that improving immune access does not necessarily eliminate immune resistance; rather, it may be accompanied by stronger IFN-γ-responsive PD-L1 expression in tumor cells, leaving PD-1/PD-L1 signaling as a potential constraint on CD8⁺ T cell activity.

Previous studies have shown that reducing stromal stiffness can increase T cell infiltration without achieving sufficient tumor control, suggesting that tumor immune infiltration and effective antitumor immune response represent distinct requirements (*9, 22*). Our findings provide a possible explanation for this apparent disconnect. Matrix softening alleviated spatial and metabolic constraints on antitumor immunity, while enhancing IFN-γ-responsive PD-L1 expression in tumor cells. This potential adaptive immune resistance may help explain the limited efficacy of matrix softening alone and provides a biological rationale for combining matrix remodeling with PD-1 blockade. The improved antitumor efficacy observed in our mouse model provides preclinical support for this combination.

Several limitations should be considered. First, the stiffness score derived from the public transcriptomic and spatial datasets reflects a stiffness-associated transcriptional state, and collagen abundance serves as an indirect histological indicator of matrix mechanics; neither provides a direct measurement of regional tissue stiffness. Future studies integrating spatial molecular profiling with regional mechanical measurements would help define this relationship more precisely. Second, the anti-PD-1 efficacy analyses were retrospective and based on relatively small cohorts, and PRJEB40416 was restricted to MSI-H GC. Larger, more diverse prospective cohorts are therefore needed to clarify the relationship between tumor matrix stiffness and response to PD-1 blockade. Third, tranilast has biological effects beyond matrix remodeling, so its *in vivo* antitumor effects cannot be attributed exclusively to changes in tumor stiffness. Complementary approaches to modifying matrix stiffness through different mechanisms should be studied further in GC to demonstrate stiffness-induced effects on tumor growth and immunotherapy response. Fourth, although LDHA knockdown attenuated both glycolytic activity and the suppressive effect of tumor cell CM on CD8⁺ T cells, the specific metabolic or secreted factors responsible for the effect on T cells remain to be identified. Finally, the molecular basis of stiffness-dependent PD-L1 induction by IFN-γ remains unresolved. Whether the increased tumor cell PD-L1 expression observed *in vivo* depends on IFN-γ and functionally limits antitumor immunity also requires direct investigation.

In conclusion, our study links matrix stiffness to spatial immune exclusion and LDHA-associated metabolic immunosuppression in GC. These findings provide a preclinical rationale for combining matrix remodeling with PD-1 blockade, and identify the accompanying PD-L1 response as a potential adaptive immune constraint which warrants further investigation.

## Materials and Methods

### Human gastric cancer specimens

Human GC tissues and paired adjacent normal tissues were obtained from 34 patients who underwent surgical resection at Xijing Hospital of Digestive Diseases, Air Force Medical University, and the Second Affiliated Hospital of Xi’an Jiaotong University. This cohort included specimens or data from our previous study (*11*) and additional patients recruited for the present study. Previously generated and new measurements were combined in the relevant analyses. Fresh tumor and matched adjacent tissues were collected after resection and allocated for mechanical measurements or histological analyses according to the experimental design. Clinicopathological variables included age, sex, histological differentiation, Lauren classification, and pathological TNM stage. Sample sizes differed among assays and are specified in the figure legends. The collection and use of human gastric cancer specimens were approved by the institutional review boards of Xijing Hospital of Digestive Diseases, Air Force Medical University (KY20192088-F-1), and The Second Affiliated Hospital of Xi ’ an Jiaotong University (2025295). Informed consent was obtained from all participants.

### Public datasets and clinical cohorts

Public datasets comprised the anti-PD-1 cohorts PRJEB25780 and PRJEB40416, the human GC scRNA-seq dataset GSE206785, the spatial transcriptomic dataset GSE270678, and The Cancer Genome Atlas Stomach Adenocarcinoma (TCGA-STAD) and Genotype-Tissue Expression (GTEx) datasets. Stiffness-associated transcriptional scores were calculated by single-sample gene set enrichment analysis (ssGSEA) using the predefined 1417-gene signature (*41*). Glycolysis scores were calculated using HALLMARK_GLYCOLYSIS gene set. Gene lists are provided in table S5. These scores represent transcriptional states associated with stiffness or glycolysis rather than direct mechanical or metabolic measurements. Computational analyses used R version 4.5.1 unless otherwise specified. Details of public dataset processing and clinical analyses are provided below.

### PRJEB25780 anti-PD-1 cohort

Publicly available bulk transcriptomic and clinical data from the PRJEB25780 gastric cancer cohort were obtained from the original study (*19*). Only tumor samples collected before pembrolizumab treatment were included. A total of 45 patients had evaluable treatment response. Treatment response was obtained from the original clinical annotations, comprising 12 responders and 33 non-responders. The tumor expression matrix was log2-transformed after addition of 1. Stiffness-associated transcriptional scores were calculated by ssGSEA using the predefined stiffness-related gene signature described above. Of the 1417 genes in the original signature, 1373 were represented in the expression dataset after gene-symbol harmonization.

Clinical PD-L1 combined positive score (CPS), microsatellite instability (MSI), and Epstein-Barr virus (EBV) status were obtained from the published clinical annotations. CPS was available for 42 of the 45 patients, and missing values were not imputed. Stiffness-associated transcriptional scores were compared between responders and non-responders using the Wilcoxon rank-sum test. The association between stiffness-associated transcriptional score and objective response was further examined using Firth logistic regression, with the stiffness-associated transcriptional score standardized within the cohort and odds ratios reported per 1-SD increase. Sensitivity analyses incorporated MSI, EBV, and continuous CPS, and leave-one-out analysis was used to assess the stability of the association. For categorical analyses, low- and high-stiffness score groups were defined using the median stiffness-associated transcriptional score of the entire cohort without data-driven optimization of the cutoff.

### PRJEB40416 anti-PD-1 cohort

Raw bulk RNA-seq data and clinical information for the PRJEB40416 cohort were obtained from the original study (*20*) and the European Nucleotide Archive. The cohort consisted of patients with advanced MSI-H gastric cancer treated with pembrolizumab. Only pretreatment tumor samples were included. Sixteen baseline samples were available for transcriptomic analysis, of which 15 had evaluable treatment response. For this cohort, objective response was defined as complete or partial response, whereas stable or progressive disease was classified as non-response, resulting in 7 responders and 8 non-responders. One patient with non-evaluable response was retained for expression processing and stiffness scoring but excluded from response analyses.

Paired-end sequencing reads were aligned to the GRCh38 primary assembly with Ensembl release 98 annotation using STAR (version 2.7.11b), and gene expression was quantified using RSEM (version 1.3.3). Gene-level TPM values below 1 were set to zero, followed by log_2_(TPM + 1) transformation. Stiffness-associated transcriptional scores were calculated by ssGSEA using the predefined stiffness-related gene signature. Scores were calculated independently within this cohort and were not derived from or normalized together with PRJEB25780.

Stiffness-associated transcriptional scores were compared between responders and non-responders. For the descriptive response analysis, low- and high-stiffness-score groups were defined using the median stiffness-associated transcriptional score. Associations with objective response were evaluated using Firth logistic regression, with stiffness-associated transcriptional scores standardized within the cohort and modeled as a continuous variable. Unadjusted and CPS-adjusted sensitivity analyses were performed in the 13 patients with available CPS data, with PD-L1 CPS entered as a continuous covariate. Leave-one-out analyses were performed to evaluate the stability of the association.

For the combined analysis of PRJEB25780 and PRJEB40416, stiffness-associated transcriptional scores were standardized within each cohort before model fitting. A Firth logistic regression model was fitted to individual patient data from both cohorts, with cohort included as a fixed effect. Odds ratios were reported per 1 SD increase in the within cohort standardized stiffness-associated transcriptional score.

### GSE206785 single-cell RNA sequencing dataset

The GSE206785 gastric cancer scRNA-seq dataset was obtained from the Gene Expression Omnibus. Analyses were restricted to 21 gastric cancer samples. Cells of low quality were excluded based on the number of detected genes, unique molecular identifier (UMI) counts, and the proportion of mitochondrial transcripts. The data were processed in Seurat with log normalization, selection of highly variable genes, principal component analysis, graph-based clustering, and UMAP visualization. Major cell populations were annotated using established lineage markers. After quality control, 99,587 cells were assigned to 10 major populations, including T/NK cells, B cells, plasma cells, myeloid cells, mast cells, endothelial cells, epithelial cells, fibroblasts, glial cells, and mural cells. The T/NK compartment was further subsetted and reclustered to identify CD4⁺ T cells, CD8⁺ T cells, and NK cells.

Stiffness-associated transcriptional scores were calculated for individual tumor samples by ssGSEA using the predefined stiffness-related gene signature. The mean stiffness-associated transcriptional score across the cohort was used to classify tumors into low- and high-stiffness score groups, comprising 15 and 6 samples, respectively. Total immune cells were defined as T/NK cells, B cells, plasma cells, myeloid cells, and mast cells. The proportions of total immune cells and CD8⁺ T cells were calculated using the total number of cells retained after quality control in each sample as the denominator. Spearman correlation analysis was used to examine the associations between stiffness-associated transcriptional score and the proportions of total immune cells and CD8⁺ T cells.

### GSE270678 spatial transcriptomic dataset

The GSE270678 gastric cancer spatial transcriptomic dataset was obtained from the Gene Expression Omnibus and comprised 72 spatial transcriptomic slices from 20 patients (*21*). Cell type deconvolution was performed using the companion scRNA-seq dataset from the same study. Tumor epithelial-enriched spots were defined by a tumor epithelial weight of at least 0.40, a tumor fraction of at least 0.80 within the epithelial compartment, a normal epithelial weight of no more than 0.15, and tumor epithelium as the dominant epithelial class. A total of 21,057 tumor epithelial-enriched spots from 70 slices were retained for downstream analyses.

Spatial gene expression counts were normalized to 10,000 counts per spot and then log transformed. Stiffness-associated transcriptional scores and glycolysis scores were calculated by ssGSEA using the predefined stiffness-related and HALLMARK_GLYCOLYSIS gene sets, respectively. Scores were calculated across the complete expression matrix of each slice and subsequently extracted for tumor epithelial-enriched spots. Within each slice, these spots were divided into low- and high-stiffness score groups according to the median stiffness-associated transcriptional score among tumor epithelial-enriched spots within that slice. LDHA positivity was defined by detectable transcript counts, whereas expression levels were evaluated using log-normalized values.

For comparisons between stiffness states, differences between high- and low-stiffness score spots were first calculated within each spatial slice. Effects from related slices that shared the same section aggregation key were averaged before further aggregation at the patient level, with patients treated as the independent units for statistical inference. Patient-level effects were evaluated using two-sided Wilcoxon signed-rank tests, and multiple comparisons were controlled using the Benjamini-Hochberg procedure.

CD8⁺ T cell abundance was represented by the CD8⁺ T cell weight obtained from spatial deconvolution. Spots with a CD8⁺ T cell weight of at least 0.05 were considered CD8-informative for the primary spatial analysis. Distances from each CD8-informative spot to the nearest high- and low-stiffness score tumor spots were calculated on the Visium hexagonal grid. The distance contrast was defined as the distance to the nearest high-stiffness score spot minus the distance to the nearest low-stiffness score spot, with positive values indicating that CD8-informative spots were farther from high-stiffness score tumor spots than from low-stiffness score tumor spots. Spatial effects were summarized at the patient level using the same hierarchical aggregation strategy.

### TCGA-STAD and GTEx datasets

Public transcriptomic data from TCGA and the GTEx datasets were analyzed using the GEPIA platform. LDHA expression was compared between gastric cancer and normal gastric tissues using log2(TPM + 1) transformed expression values. Normal controls included available TCGA normal tissues together with normal gastric tissues from GTEx. The relationship between YAP1 and LDHA expression was assessed in TCGA STAD tumor samples using Spearman correlation analysis.

### Cell culture and hydrogel preparation

AGS and MKN45 human GC cells were kindly provided by the State Key Laboratory of Cancer Biology, Xijing Hospital. MFC cells were purchased from Procell Life Science & Technology. All cell lines were authenticated and tested negative for mycoplasma contamination. Cells were cultured in RPMI 1640 containing 10% fetal bovine serum and 1% penicillin/streptomycin at 37°C in a humidified atmosphere containing 5% CO₂. PA hydrogels of different stiffness were prepared by varying the concentrations of acrylamide and bis-acrylamide. Hydrogels with storage moduli (G′) of approximately 0.2 and 6.4 kPa were used as soft and stiff culture substrates, respectively, based on our previous study (*11*). For cell attachment, the hydrogel surfaces were functionalized with sulfo-SANPAH under UV irradiation and subsequently coated with type I collagen. Cells were seeded on the hydrogels and cultured for subsequent experiments.

### Mouse tumor model and treatment

Female 615 mice aged 6–8 weeks were used and housed under specific pathogen-free conditions. A syngeneic subcutaneous gastric tumor model was established on day 0 by subcutaneous inoculation of MFC cells (3 × 10⁶ cells in 0.1 mL of suspension per mouse) into the left lower abdomen. Mice were randomly assigned to control, tranilast, anti-PD-1, or tranilast plus anti-PD-1 treatment groups. The experimental design, including tumor implantation and treatment schedule, is shown in Fig. 2A. Tranilast was prepared in 1% sodium carboxymethylcellulose (CMC-Na) and administered by oral gavage at 200 mg/kg daily on days 10–20. Anti-mouse PD-1 antibody was administered intraperitoneally at 10 mg/kg on days 12, 15, and 18. The combination group received both regimens, whereas control mice received CMC-Na by oral gavage and PBS by intraperitoneal injection.

Two separate cohorts were used: the first comprised five mice per group for endpoint tissue analyses, and the second comprised three mice per group for ¹⁸F-FDG PET/CT. Tumor growth was monitored throughout the experiment in both cohorts. Tumor length and width were measured every 2–3 days. Tumor volume (mm³) was calculated as length (mm) × width (mm)²/2. At the experimental endpoint, mice in the first cohort were euthanized by cervical dislocation. Tumors were excised, weighed, and collected for mechanical, histological, immunological, and transcriptomic analyses. For scRNA-seq, tumors from two mice per group were processed individually. Major organs were collected for histopathological assessment. Mice in the second cohort underwent ¹⁸F-FDG PET/CT at the experimental endpoint to assess tumor glucose uptake, quantified as the maximum standardized uptake value (SUVmax). After imaging, mice were euthanized by cervical dislocation, and subcutaneous tumors were excised for measurement of tumor lactate levels. Assay-specific sample sizes are provided in the figure legends. All animal experiments were approved by the Institutional Animal Care and Use Committee of Xi’an Jiaotong University (XJTUAE2025-2673).

### Mechanical measurements

Mechanical properties of human gastric tissues were assessed by rotational rheometry, and atomic force microscopy (AFM); mouse tumors were assessed by rotational rheometry. For rheological measurements, fresh tissue specimens were tested at 37°C and the storage modulus (G′) was recorded. For AFM measurements, frozen human tissue sections were indented in PBS, and force-indentation curves were fitted using the Hertz model with a Poisson ratio of 0.5 to calculate the apparent Young’s modulus. For integrated analysis of human gastric tissues, rheometry-derived storage modulus values were converted to estimated Young’s modulus according to E = 2G′(1+ν), with ν set to 0.5. This conversion treats the measured storage modulus as an elastic shear-modulus approximation; converted values are estimates rather than independent AFM measurements. Multiple measurements from each specimen were averaged for subsequent analysis. The mechanical properties of mouse tumors and PA hydrogels are reported as storage modulus (G′).

### Histology and tissue fluorescence imaging

Formalin-fixed, paraffin-embedded human and mouse tissues were analyzed by hematoxylin and eosin, Masson’s trichrome, Sirius red, immunohistochemical, and immunofluorescence staining. SHG imaging was performed on H&E-stained sections using multiphoton microscopy. Multiplex immunofluorescence assays were performed using the commercial multiplex fluorescence staining kit according to the manufacturer’s instructions. Images were acquired using a FLUOVIEW FV4000 confocal laser scanning microscope (Evident, formerly Olympus) or an Akoya multispectral fluorescence imaging system, as appropriate. Antibodies are listed in table S6. Quantitative image analysis was performed to assess staining area, fluorescence intensity, positive cell abundance, subcellular localization, and spatial relationships between defined cell populations. For within-patient regional analyses of human GC tissues, collagen I-low and collagen I-high areas were used as histological proxies for relatively lower and higher matrix stiffness. Matched regions on serial sections were used where indicated; regional collagen abundance was not treated as a direct stiffness measurement. Patient-level values summarized the corresponding sampled fields. GC specimens were classified into low- and high-stiffness groups according to the median measured tissue stiffness.

### Cellular fluorescence imaging

YAP localization and LDHA expression were assessed by immunofluorescence in GC cells on soft or stiff PA hydrogels, with the indicated drug treatments. Nuclei were counterstained with DAPI. YAP localization was quantified as the nuclear-to-cytoplasmic fluorescence intensity ratio, and LDHA expression as mean fluorescence intensity (MFI). Phalloidin staining assessed F-actin organization, with cell area and circularity quantified from cell outlines. Antibodies are listed in table S6.

### RNA sequencing and bioinformatic analyses

Single-cell transcriptomic profiling was performed on fresh mouse tumors collected from the control, tranilast, anti-PD-1, and tranilast combined with anti-PD-1 groups (n = 2 mice/group). Each tumor was obtained from a separate mouse and processed as an individual biological sample. Tumors were enzymatically dissociated, and viable single-cell suspensions were processed on the 10X Genomics Chromium platform. Sequencing reads were processed with Cell Ranger. Low-quality cells were excluded based on detected gene numbers, UMI counts, and mitochondrial transcript proportions. Data were processed in Seurat (version 4.0.3) using log normalization, highly variable gene selection, principal component analysis, graph-based clustering, and UMAP visualization. Major cellular compartments were defined according to canonical lineage markers. Tumor epithelial cells and CD8⁺ T cells were subsequently analyzed at higher resolution to resolve transcriptionally distinct subpopulations.

Cell-level differential expression analysis was performed using the FindMarkers function with the Wilcoxon rank-sum test. DEGs were selected using thresholds of |log₂ FC| > 0.25 and adjusted *P* < 0.05, with a minimum detection fraction of 0.1. Gene Ontology biological process enrichment analysis was performed using clusterProfiler (version 4.6.2). QuSAGE was applied to the expression matrices of tumor epithelial cells to compare metabolic pathway activity across the four treatment groups.

Seven CD8⁺ T cell subclusters were grouped into memory/progenitor-like, intermediate/proliferative, and terminal/ISG-associated states. Epithelial reclustering identified Epi-1–Epi-5 and a PD-L1-high epithelial state. Displayed subcluster proportions used the corresponding parent population in each treatment group as the denominator.

Bulk RNA-seq was performed on AGS cells cultured on soft and stiff PA hydrogels for 48 h (n = 3). DEGs between the two stiffness conditions were identified using thresholds of |log₂ FC| > 0.25 and adjusted *P* < 0.05, and GSEA was performed to characterize biological processes associated with matrix stiffness. Gene sets related to glycolysis, YAP/TAZ, and actin cytoskeleton reorganization were examined. The bulk RNA-seq results were compared with DEGs in tumor epithelial cells from the mouse single-cell dataset in the tranilast versus control and tranilast plus anti-PD-1 versus anti-PD-1 comparisons to identify shared stiffness-associated genes.

### Metabolic assays

Glucose uptake was measured using the fluorescent glucose analogue 2-NBDG. Glucose and lactate concentrations were measured in culture medium and cell lysates to determine extracellular and intracellular levels, respectively. Lactate levels were also measured in mouse tumor tissues. LDH activity was measured in cell lysates. Intracellular glucose and lactate levels, tumor lactate levels, and cellular LDH activity were normalized to the protein concentrations of the corresponding lysates. All assays were performed according to the manufacturers’ instructions.

### Conditioned medium experiments and flow cytometry

CM was collected from MFC cells cultured on soft or stiff PA hydrogels and used to culture mouse splenic lymphocytes. For LDHA knockdown experiments, CM was collected from control or LDHA-knockdown MFC cells cultured on stiff PA hydrogels. Splenic lymphocytes isolated from healthy mice were pooled and expanded in culture before CM treatment. After 24 h of CM treatment, the proportions of PD-1⁺TIM-3⁺ and IFN-γ⁺TNF-α⁺ cells within the CD8⁺ T cell population were quantified by flow cytometry.

In separate experiments, GC cells cultured on soft or stiff PA hydrogels were stimulated with IFN-γ (100 ng/mL) for 24 h, and surface expression of PD-L1 and MHC/HLA molecules was analyzed by flow cytometry. Surface expression was quantified as the percentage of positive cells and median fluorescence intensity (MFI). For each cell line and hydrogel stiffness, MFI fold change was calculated as stimulated MFI/unstimulated MFI, and ΔMFI as stimulated MFI − unstimulated MFI. Both measures were calculated separately for each independent experiment using the corresponding unstimulated control. Both the CM treatment experiments and the IFN-γ stimulation experiments were performed in three independent experiments. Flow cytometry data were analyzed using FlowJo software.

### Genetic and pharmacological interventions

LDHA and YAP expression was reduced by RNA interference. The RNA interference reagents used in this study are listed in table S6. FX11 (5 μM, 24 h) and verteporfin (2 μM, 24 h) were used to inhibit LDHA enzymatic activity and YAP transcriptional activity, respectively, whereas latrunculin A (Lat.A, 0.2 μM, 24 h) was used to disrupt actin polymerization.

### Western blotting and ChIP-qPCR

Proteins were extracted using RIPA lysis buffer. Western blotting was performed using standard procedures. Protein bands were visualized using enhanced chemiluminescence and quantified by densitometry using ImageJ. β-actin was used as the loading control for normalization. The p-YAP/total YAP ratio was calculated by dividing the p-YAP band intensity by the total YAP band intensity. ChIP-qPCR was performed in AGS cells treated with vehicle or verteporfin using a commercial ChIP kit and an antibody against YAP, with IgG used as a negative control. Enrichment of the indicated LDHA promoter region was quantified relative to input DNA and expressed as a percentage of input. Antibody, primer, and kit information is provided in table S6.

### Statistical analysis

Quantitative data are presented as mean ± SD unless otherwise indicated. Statistical analyses were performed using GraphPad Prism (version 8.0) and R (version 4.5.1). Two-group comparisons were performed using paired or unpaired t tests, Mann-Whitney U tests, or Wilcoxon signed-rank tests, as appropriate. Comparisons among multiple groups were performed using one-way analysis of variance (ANOVA). Spearman rank correlation was used for correlation analyses, and Firth logistic regression was applied to assess the association between stiffness-associated transcriptional score and response to PD-1 blockade. Spatial analyses used patient-level estimates, with Benjamini–Hochberg correction for multiple testing. Treatment interactions were assessed using two-way ANOVA and eGR-based Bliss analysis. Tests were two-sided except for the one-sided bootstrap tests used in eGR-based Bliss analysis. *P* value < 0.05 was considered statistically significant.

## Abbreviations

AFM: atomic force microscopy
AUC: area under the curve
bulk RNA-seq: bulk RNA sequencing
ChIP-qPCR: chromatin immunoprecipitation followed by quantitative PCR
CI: confidence interval
CM: conditioned medium
CPS: combined positive score
DEGs: differentially expressed genes
EBV: Epstein–Barr virus
ECM: extracellular matrix
eGR: exponential growth rate
GC: gastric cancer
GSEA: gene set enrichment analysis
GTEx: Genotype-Tissue Expression
H&E: hematoxylin and eosin; ISG, interferon-stimulated gene
Lat.A: latrunculin A
LDH: lactate dehydrogenase
MFI: mean fluorescence intensity (microscopy) or median fluorescence intensity (flow cytometry)
MSI: microsatellite instability
MSI-H: microsatellite instability-high
OR: odds ratio
PA: polyacrylamide
PanCK: pan-cytokeratin
QuSAGE: quantitative set analysis for gene expression
ROC: receiver operating characteristic
scRNA-seq: single-cell RNA sequencing
SHG: second harmonic generation
ssGSEA: single-sample gene set enrichment analysis
SUVmax: maximum standardized uptake value
TCGA-STAD: The Cancer Genome Atlas Stomach Adenocarcinoma
Tcm: central memory T cells
TEX: exhausted T cells
UMAP: uniform manifold approximation and projection
UMI: unique molecular identifier
VP: verteporfin

## Acknowledgements

We thank the patients who contributed tissue samples to this study. We also thank the State Key Laboratory of Cancer Biology, Xijing Hospital, for providing the AGS and MKN45 cell lines.

## Funding

This work was supported by the National Natural Science Foundation of China (No. 82303812, No. 82403890 and No. 82560549; Key Program, No. 12432015) and Joint Founding Project of Innovation Research Institute, Xijing Hospital (LHJJ24JH28).

## Author contributions

Q.S., J. Wu, Z.W., and Y.C. contributed equally to this work. J. Wu and N.X. conceived and designed the study. Q.S. and Z.W. established the experimental models and performed the major experiments. Y.C., X.L., H.T., Y.P., B.Q., and B.Z. collected clinical specimens and curated the clinical data. Q.S., Y.C., and X.L. performed the bioinformatic and statistical analyses. Y.L. and H.S. analyzed and interpreted the experimental data. Q.S., J. Wu, Z.W., and X.L. prepared the figures and visualizations. Q.S., J. Wu, and N.X. drafted the manuscript. F.X., J. Wang, Y.N., Y.S., N.L., and D.F. critically revised the manuscript for important intellectual content. J. Wu, F.X., N.L., and N.X. acquired funding. T.W., L.L., and N.X. supervised the study. All authors reviewed and approved the final version of the manuscript.

## Conflict of interest

The authors declare no competing interests.

## Data availability

The scRNA-seq data from mouse tumors has been deposited in the National Genomics Data Center (GSA; accession number: PRJCA077272). The bulk RNA-seq data from AGS cells has been deposited in the National Genomics Data Center (GSA for Human; accession number: PRJCA077376).

## Notes

### Competing Interest Statement

The authors have declared no competing interest.

