## Supplemental figures 1-11 for "Matrix stiffness shapes immunometabolic states and response to PD-1 blockade in gastric cancer"

**Supplementary figures with legends**

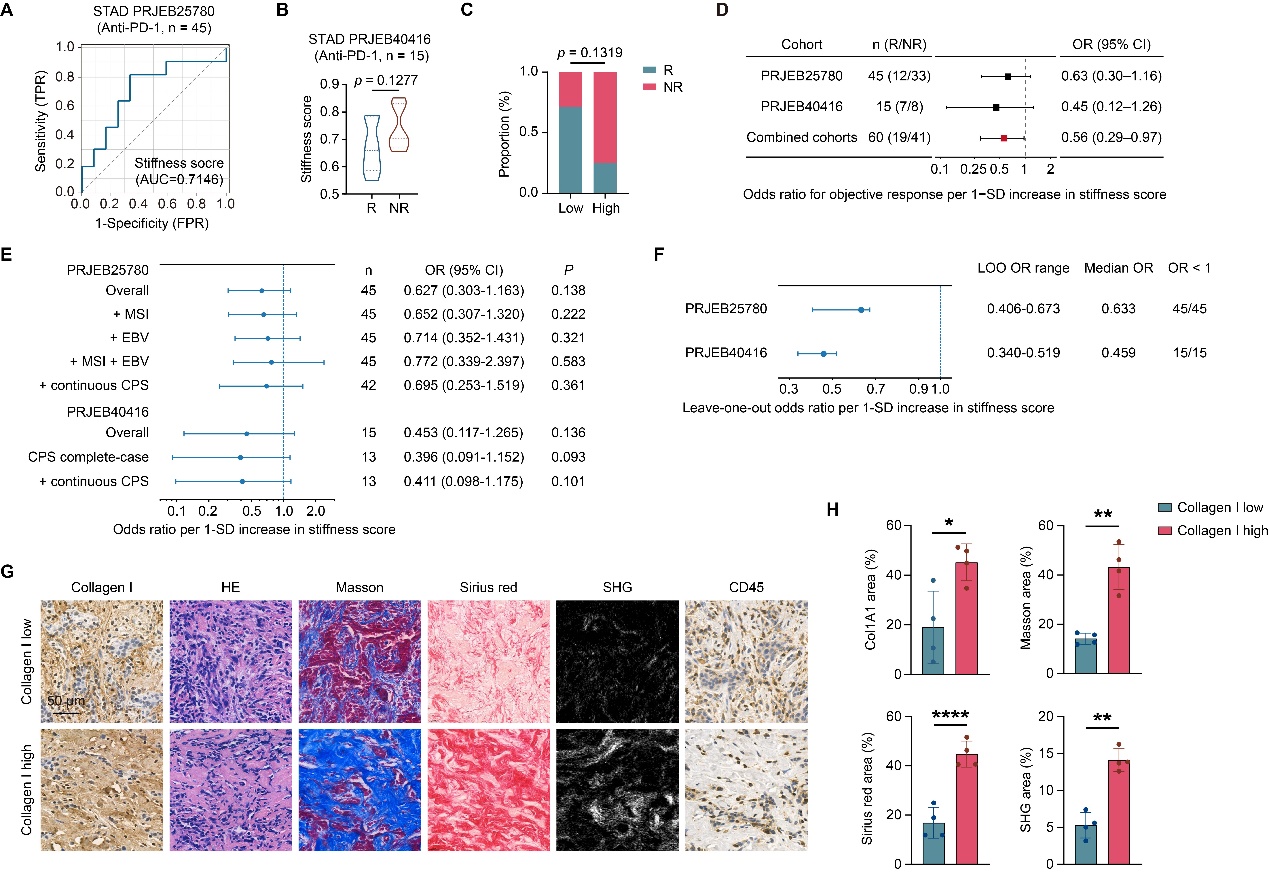

**Fig. S1. Matrix stiffness is associated with anti-PD-1 response and immune infiltration in gastric cancer.** (**A**) Receiver operating characteristic (ROC) curve for the stiffness-associated transcriptional score in the PRJEB25780 anti-PD-1 cohort. (**B**) Stiffness-associated transcriptional scores in responders (R; n = 7) and non-responders (NR; n = 8) in PRJEB40416. Lines indicate the median and quartiles. (**C**) Response distributions in low- and high-stiffness-score groups in PRJEB40416, according to the median stiffness-associated transcriptional score (n = 7 and 8, respectively). (**D**) Cohort-specific and combined associations between stiffness-associated transcriptional scores and objective response estimated by Firth logistic regression. The combined model includes cohort as a fixed effect. Black and red squares indicate cohort-specific and combined odds ratios (ORs), respectively; bars show 95% confidence intervals (CIs). (**E**) Sensitivity analyses adjusted for microsatellite instability (MSI), Epstein-Barr virus (EBV), both, or continuous PD-L1 combined positive score (CPS) in PRJEB25780, and unadjusted and CPS-adjusted analyses in the CPS-complete subset of PRJEB40416. Unadjusted full-cohort estimates are shown for comparison. (**F**) Leave-one-out Firth regression analyses in PRJEB25780 (45 iterations) and PRJEB40416 (15 iterations). Points indicate median ORs; bars indicate minimum-to-maximum ranges, not 95% CIs. ORs in (**D** to **F**) are per 1-SD increase in the within-cohort standardized score; dashed lines indicate OR = 1. (**G** and **H**) Representative collagen I and CD45 immunohistochemistry, H&E, Masson’s trichrome, Sirius red, and SHG images (**G**), with paired quantification of positive areas for collagen I, Masson’s trichrome, Sirius red, and SHG (**H**), in collagen I-low/high regions from serial sections of human GC specimens (n = 4 patients; one point per patient). Scale bars, 50 μm. **P* < 0.05, ***P* < 0.01, *****P* < 0.0001.

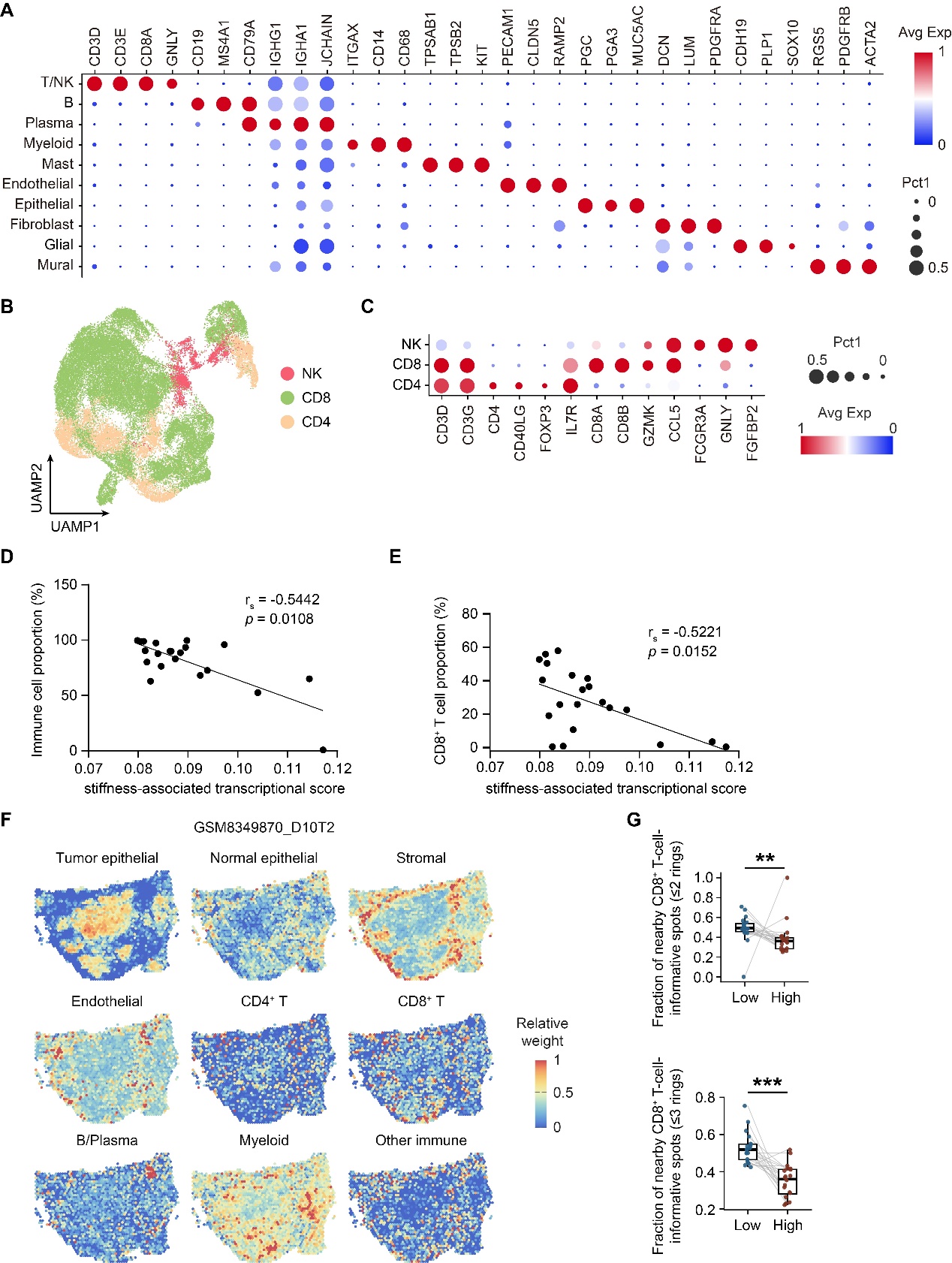

**Fig. S2. Immune composition and spatial distribution associated with stiffness scores in gastric cancer.** (**A**) Dot plot of marker expression for the 10 major cell populations in GSE206785. (**B**) Uniform manifold approximation and projection (UMAP) of reclustered T/NK cells. (**C**) Dot plot of marker expression for T/NK cell subpopulations. Dot size indicates the expressing cell fraction; color indicates scaled average expression in (**A** and **C**). (**D** and **E**) Spearman correlations of stiffness-associated transcriptional scores with total immune cell (**D**) and CD8⁺ T cell (**E**) proportions (n = 21). Total immune cells include T/NK cells, B cells, plasma cells, myeloid cells, and mast cells. (**F**) Spatial maps of relative deconvolution weights for nine major cell populations in GC section GSM8349870_D10T2. (**G**) Paired patient-level comparisons of the fractions of CD8⁺ T cell-informative spots located within two spot rings (top) or three spot rings (bottom) of low- and high-stiffness-score tumor spots (n = 20). ***P* < 0.01, ****P* < 0.001.

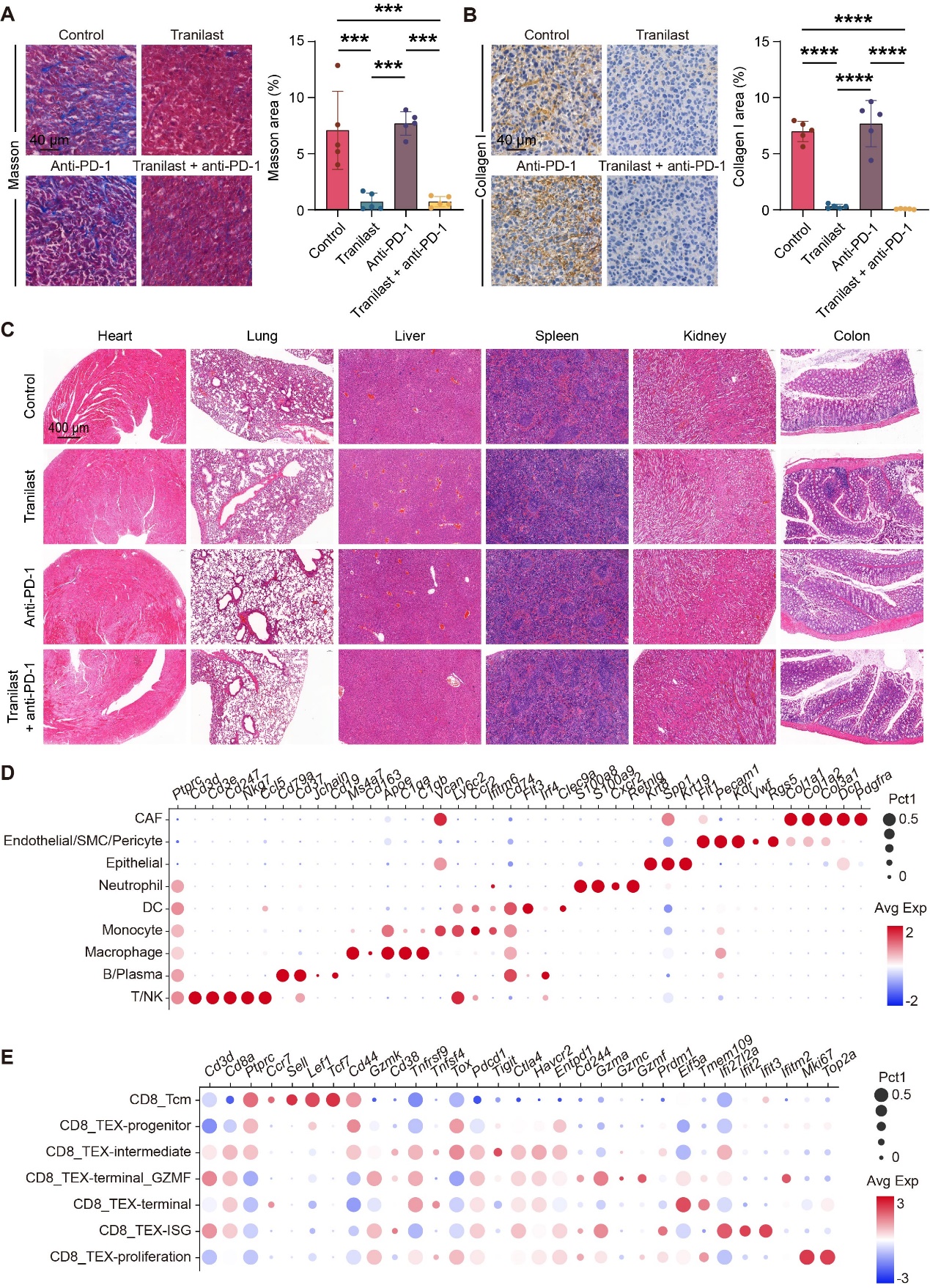

**Fig. S3. Matrix remodeling, organ histology, and single-cell annotation in the 615 mouse tumor model.** (**A** and **B**) Representative Masson’s trichrome staining (**A**) and collagen I immunohistochemistry (**B**), with quantification of positive areas in subcutaneous tumors from the indicated treatment groups (n = 5 mice/group; one point/mouse). Scale bars, 40 μm. (**C**) Representative hematoxylin and eosin (H&E) staining of major organs from mice in the indicated treatment groups. Scale bars, 400 μm. (**D** and **E**) Dot plots of marker expression for major cell populations (**D**) and seven CD8⁺ T cell subclusters (**E**). Dot size indicates the expressing cell fraction; color indicates scaled average expression. CD8⁺ T cell subclusters were grouped into memory/progenitor-like, intermediate/proliferative, and terminal/interferon-stimulated gene (ISG)-associated states in **Fig. 2K**. ****P* < 0.001, *****P* < 0.0001.

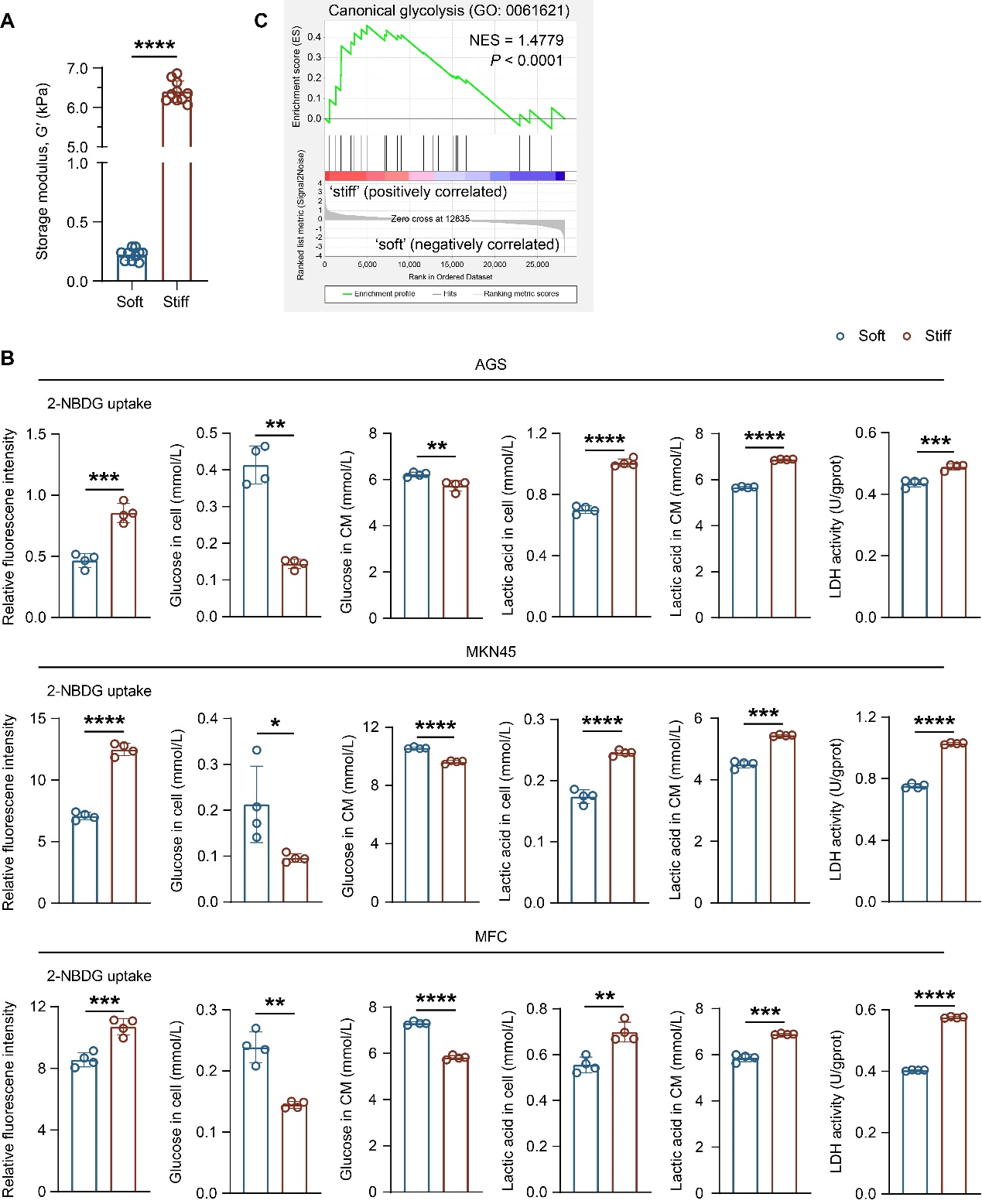

**Fig. S4. Matrix stiffness promotes glycolytic reprogramming in gastric cancer cells.** (**A**) Storage modulus (G′) of soft and stiff polyacrylamide (PA) hydrogels measured by rotational rheometry (N ≥ 3 independent experiments, n = 10 measurements/group). (**B**) 2-NBDG uptake, intracellular glucose (Glu-cell), residual medium glucose (Glu-CM), intracellular lactate (Lac-cell), medium lactate (Lac-CM), and cellular LDH activity in AGS, MKN45, and MFC cells on soft or stiff PA hydrogels (n = 4). (**C**) Gene set enrichment analysis (GSEA) of canonical glycolysis in bulk RNA-seq data from AGS cells on stiff versus soft PA hydrogels. **P* < 0.05, ***P* < 0.01, ****P* < 0.001, *****P* < 0.0001.

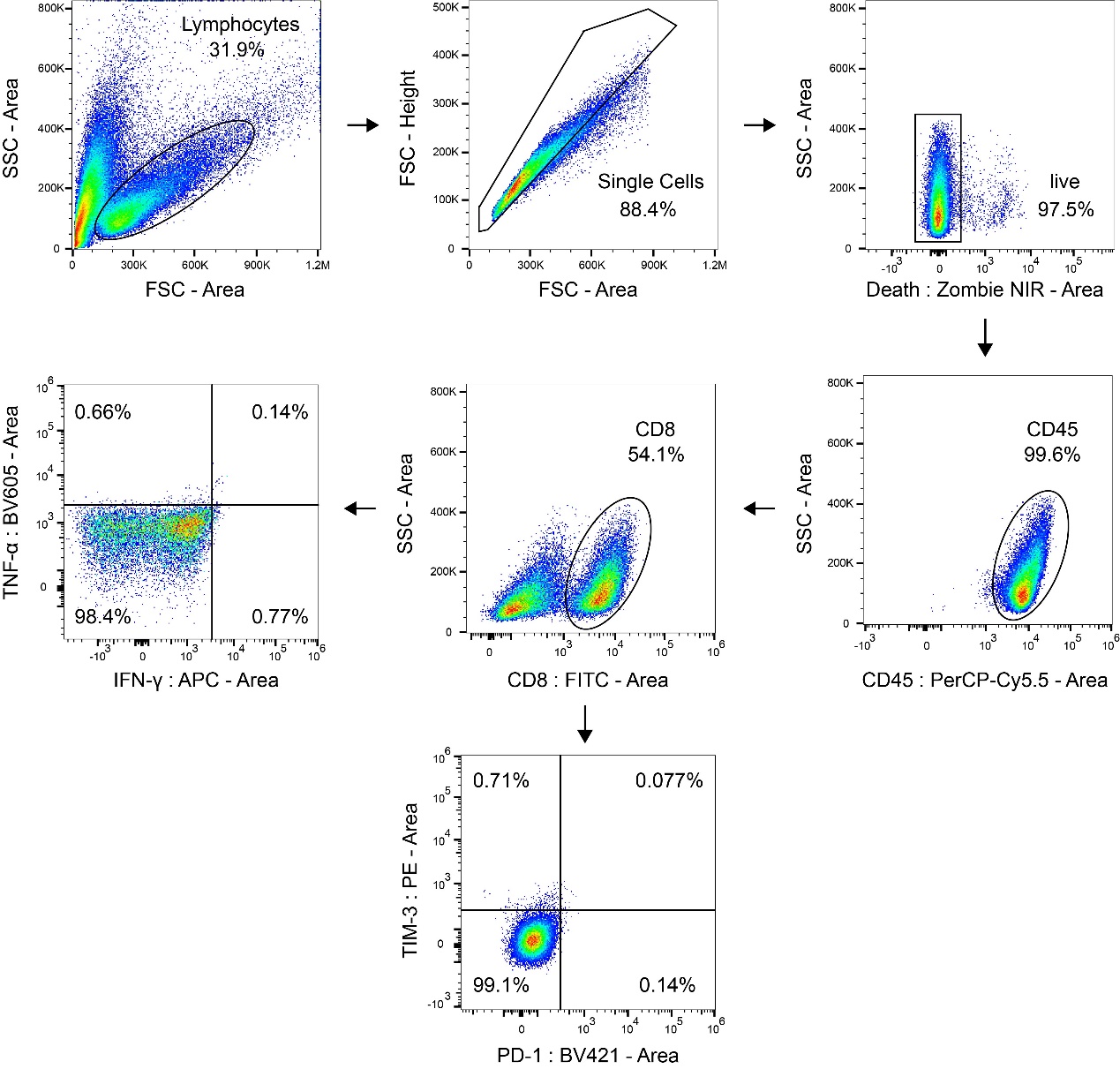

**Fig. S5. Flow cytometry gating strategies for CD8⁺ T cell analyses.** Representative gating for identification of CD8⁺ T cells and quantification of IFN-γ⁺TNF-α⁺ and PD-1⁺TIM-3⁺ fractions within the CD8⁺ T cell population.

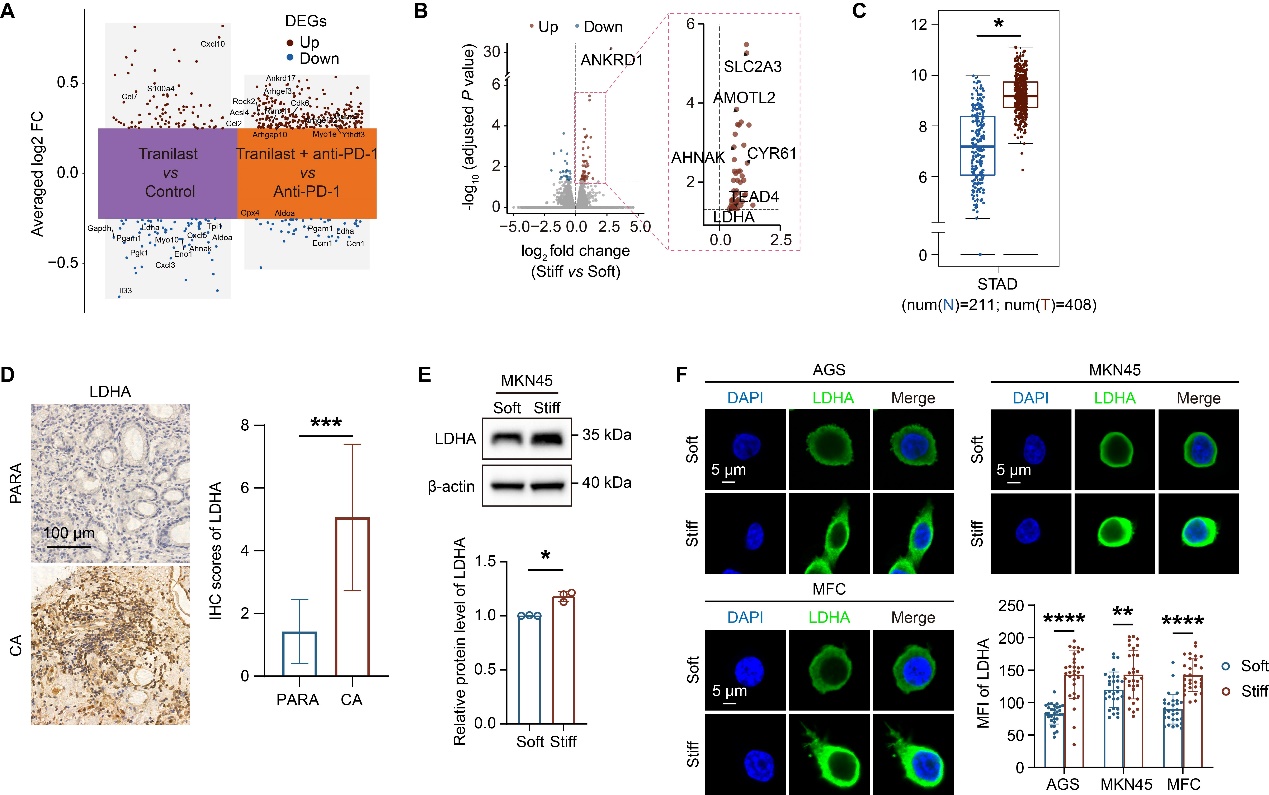

**Fig. S6. Matrix stiffness is associated with LDHA expression in gastric cancer.** (**A**) Volcano plots of differentially expressed genes (DEGs) in tumor epithelial cells from mouse scRNA-seq in the tranilast versus control and combination versus anti-PD-1 comparisons. DEG thresholds are defined in the Methods. (**B**) Volcano plot of DEGs in AGS cells on stiff versus soft PA hydrogels, highlighting ANKRD1, AMOTL2, CYR61, and LDHA. (**C**) LDHA mRNA expression in human GC and normal gastric tissues from public transcriptomic datasets. (**D**) Representative LDHA immunohistochemistry and paired quantification in human GC and matched adjacent normal tissues (n = 11). Scale bars, 100 μm. (**E**) Representative LDHA Western blots and quantification in MKN45 cells on soft or stiff PA hydrogels (n = 3). (**F**) Representative LDHA immunofluorescence and mean fluorescence intensity in AGS, MKN45, and MFC cells on soft or stiff PA hydrogels (N = 3 independent experiments, n = 30 cells/condition). Scale bars, 5 μm. **P* < 0.05, ***P* < 0.01, ****P* < 0.001, *****P* < 0.0001.

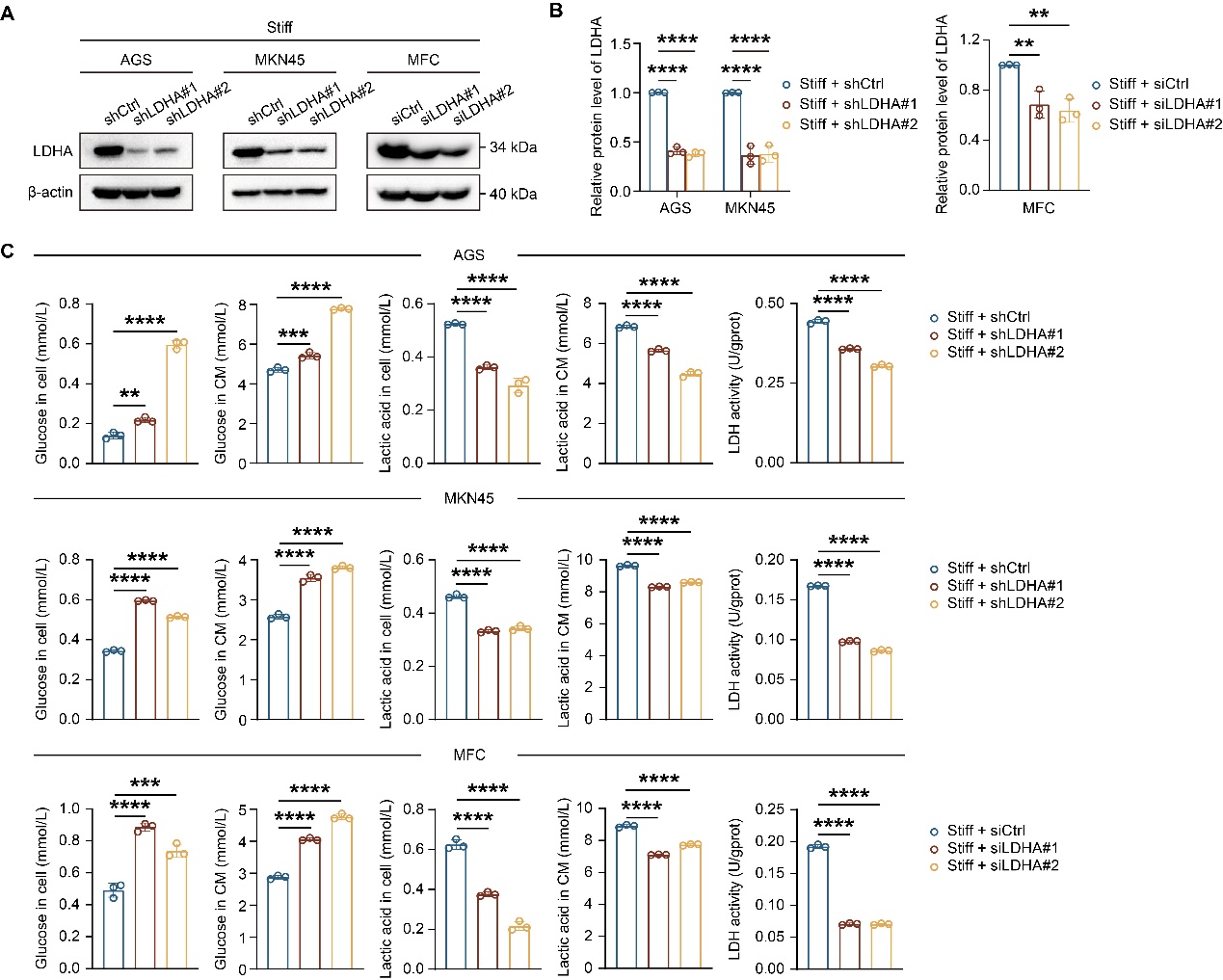

**Fig. S7. LDHA knockdown attenuates glycolytic reprogramming induced by high matrix stiffness in gastric cancer cells.** (**A** and **B**) Representative LDHA Western blots (**A**) and knockdown efficiency (**B**) in AGS, MKN45, and MFC cells on stiff PA hydrogels (n = 3). (**C**) Intracellular glucose, residual medium glucose, intracellular lactate, medium lactate, and cellular LDH activity after LDHA knockdown on stiff PA hydrogels (n = 3). ***P* < 0.01, ****P* < 0.001, *****P* < 0.0001.

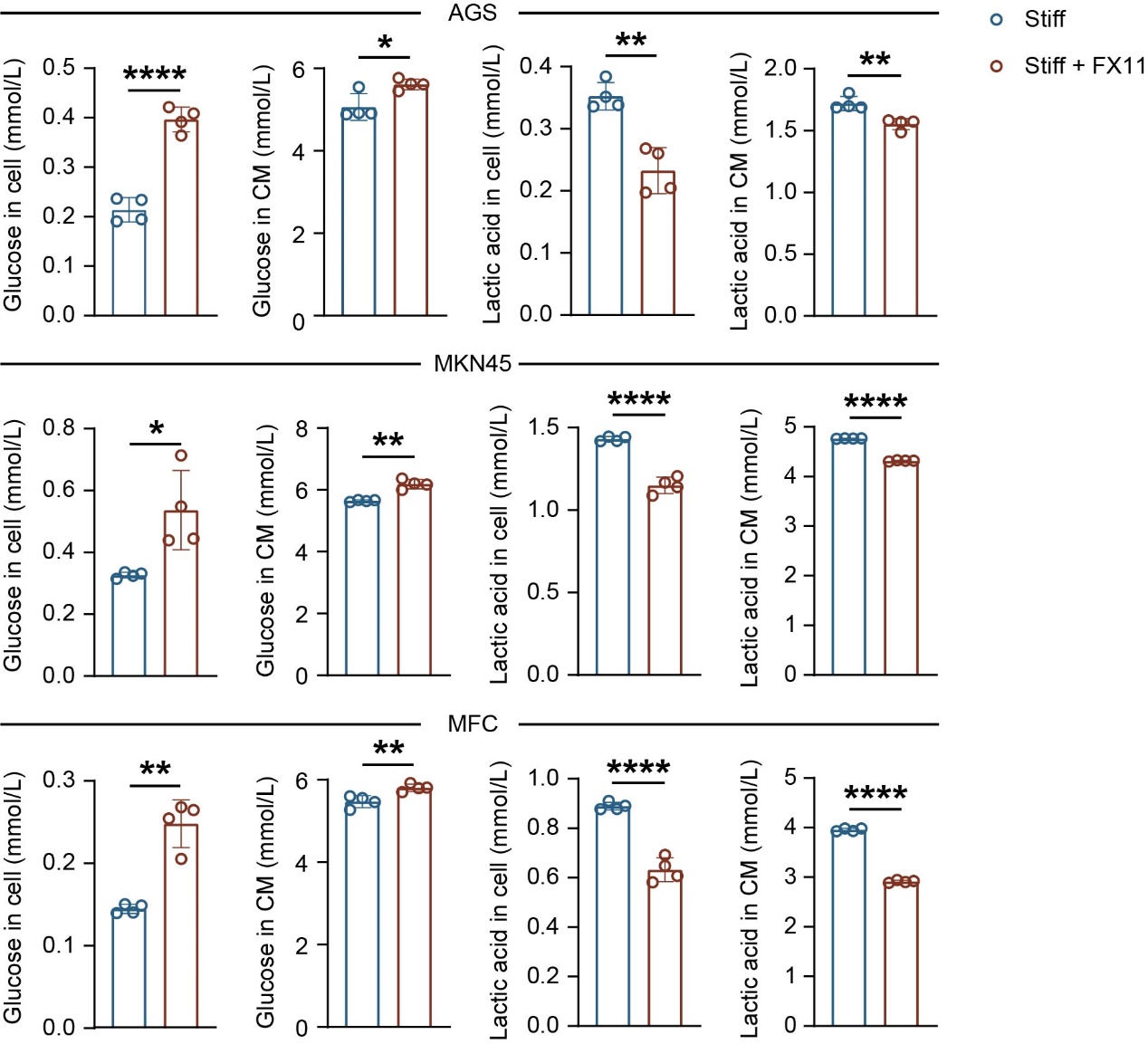

**Fig. S8. Pharmacological inhibition of LDHA attenuates glycolytic reprogramming induced by high matrix stiffness in gastric cancer cells.** Intracellular glucose, residual medium glucose, intracellular lactate, and medium lactate in AGS, MKN45, and MFC cells on stiff PA hydrogels with vehicle or the LDHA inhibitor FX11 (n = 4). **P* < 0.05, ***P* < 0.01, *****P* < 0.0001.

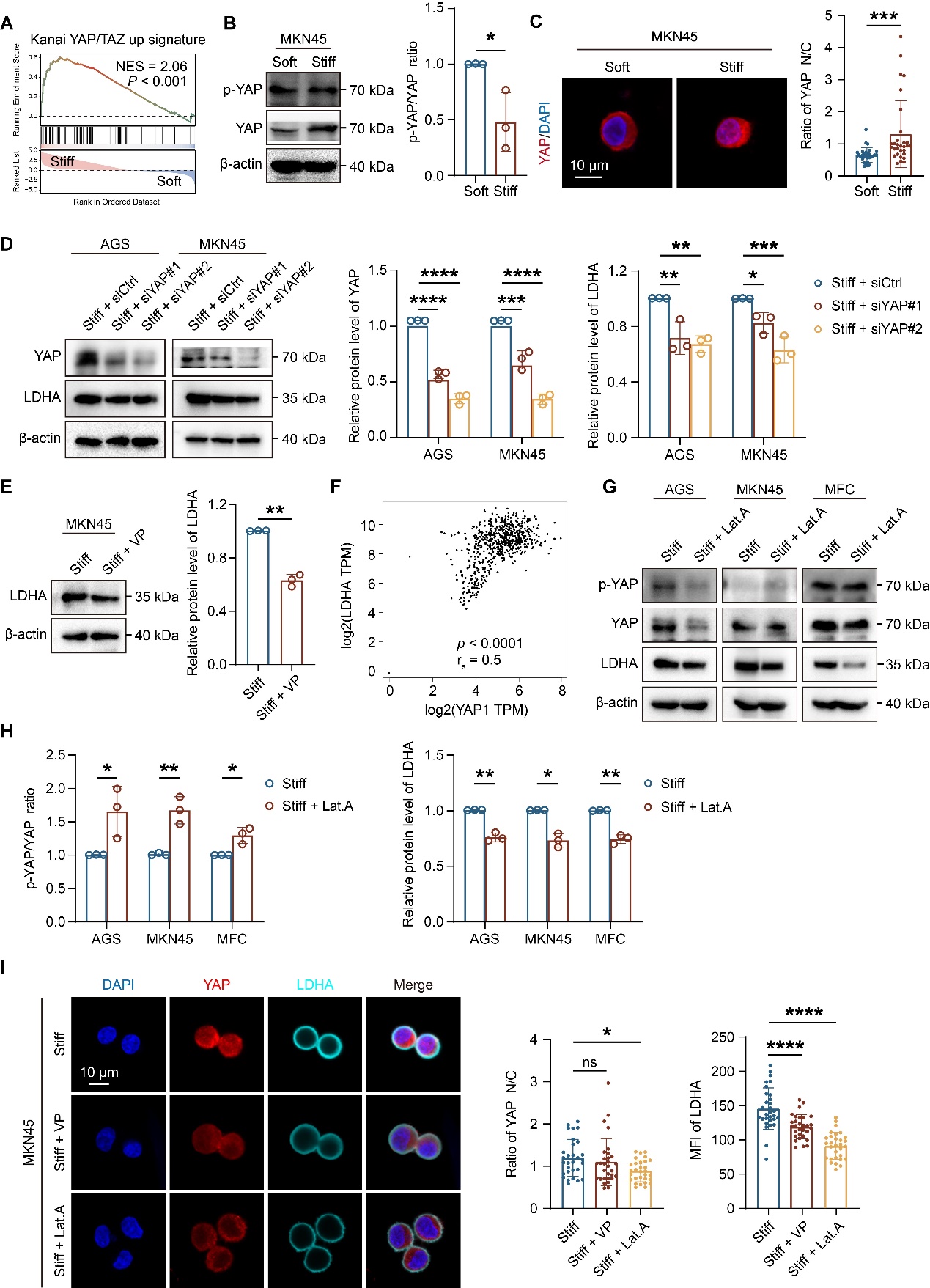

**Fig. S9. F-actin and YAP mediate stiffness-dependent LDHA expression in gastric cancer cells.** (**A**) Gene set enrichment analysis (GSEA) of the Kanai YAP/TAZ up signature in bulk RNA-seq data from AGS cells on stiff versus soft PA hydrogels. (**B**) Representative Western blots of phosphorylated YAP (p-YAP) and total YAP, with p-YAP/YAP quantification in MKN45 GC cells on soft or stiff PA hydrogels (n = 3). (**C**) Representative YAP immunofluorescence and nuclear-to-cytoplasmic intensity ratios in MKN45 cells on soft or stiff PA hydrogels (N = 3 independent experiments, n = 30 cells/condition). Scale bars, 10 μm. (**D**) YAP knockdown validation and LDHA protein expression in AGS and MKN45 cells on stiff PA hydrogels assessed by Western blotting (n = 3). (**E**) Representative LDHA Western blots and quantification in MKN45 cells on stiff PA hydrogels with vehicle or verteporfin (VP) (n = 3). (**F**) Spearman correlations between YAP1 and LDHA mRNA expression in human GC samples from public transcriptomic datasets. (**G** and **H**) Representative p-YAP, total YAP, and LDHA Western blots (**G**), with p-YAP/YAP ratios and relative LDHA protein expression (**H**), in AGS, MKN45, and MFC cells on stiff PA hydrogels with vehicle or latrunculin A (Lat.A) (n = 3). (**I**) Representative YAP and LDHA immunofluorescence, YAP nuclear-to-cytoplasmic intensity ratios, and LDHA mean fluorescence intensity in MKN45 cells on stiff PA hydrogels with vehicle, VP, or Lat.A (N = 3 independent experiments, n = 30 cells/condition). Scale bars, 10 μm. ns, *P* > 0.05, **P* < 0.05, ***P* < 0.01, ****P* < 0.001, *****P* < 0.0001.

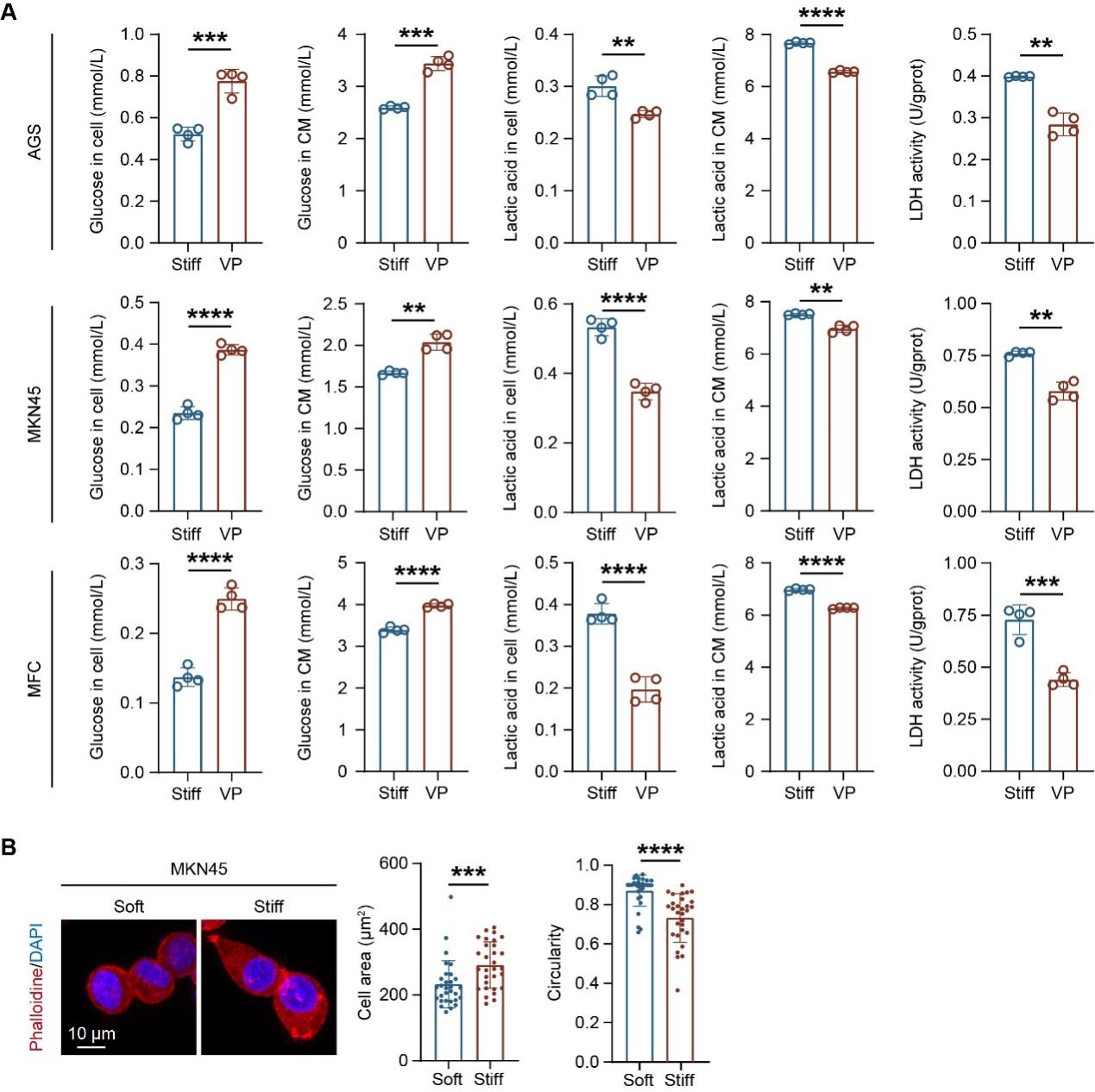

**Fig. S10. YAP-dependent glycolysis and stiffness-associated F-actin organization in gastric cancer cells.** (**A**) Intracellular glucose, residual medium glucose, intracellular lactate, medium lactate, and cellular LDH activity in AGS, MKN45, and MFC cells on stiff PA hydrogels with vehicle or verteporfin (VP) (n = 4). (**B**) Representative phalloidin staining and quantification of cell area and circularity in MKN45 cells on soft or stiff PA hydrogels (N = 3 independent experiments, n = 30 cells/condition). Scale bars, 10 μm. ***P* < 0.01, ****P* < 0.001, *****P* < 0.0001.

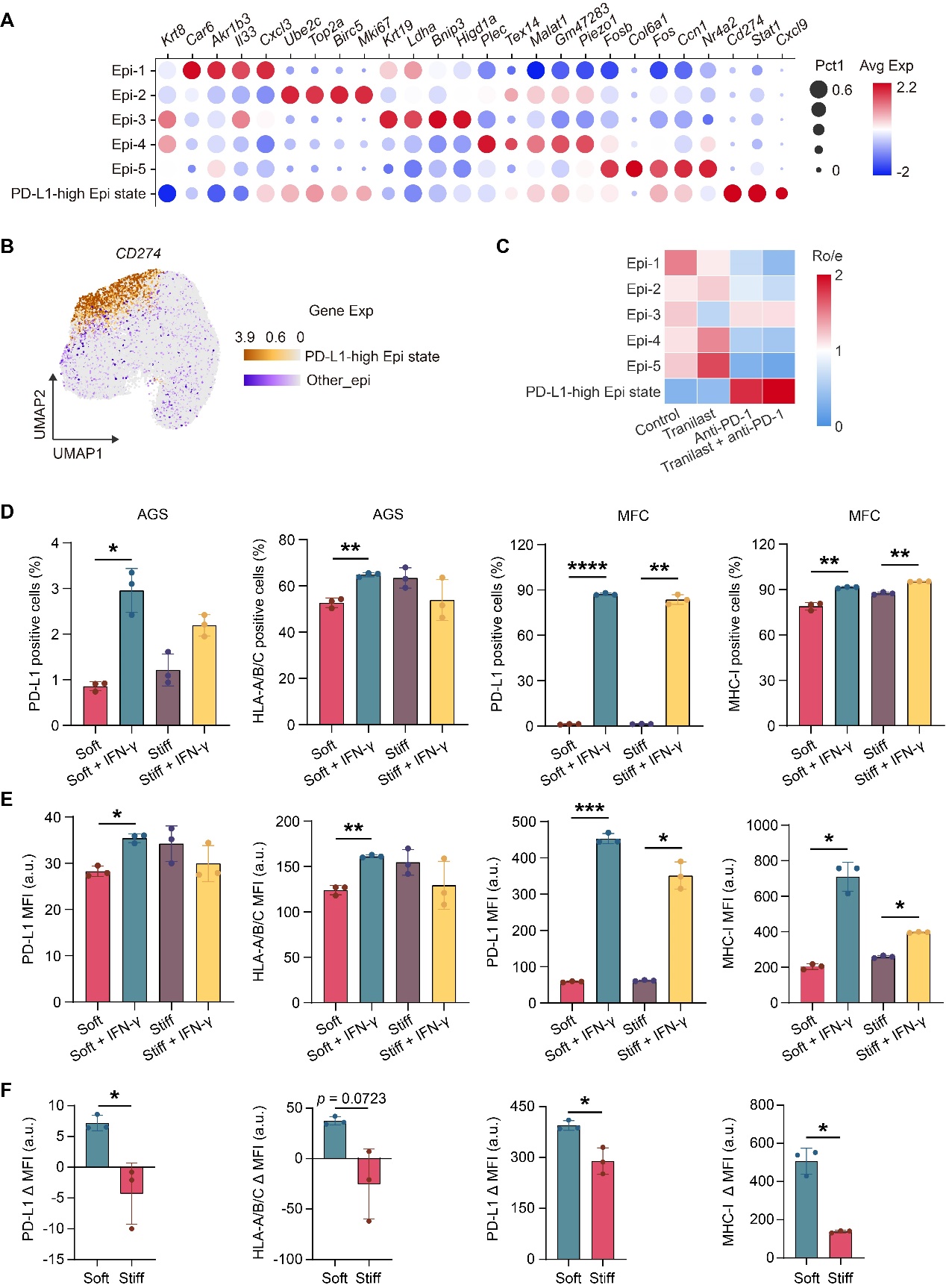

**Fig. S11. The PD-L1-high epithelial state and stiffness-dependent IFN-γ responses.** (**A**) Dot plot of marker expression for epithelial subclusters in scRNA-seq of 615 mouse subcutaneous tumors. Dot size indicates the expressing cell fraction; color indicates scaled average expression. (**B**) UMAP of *Cd274* expression highlighting the PD-L1-high epithelial state relative to the remaining epithelial cells. (**C**) Relative abundance of epithelial subclusters across control, tranilast, anti-PD-1, and combination groups, expressed as observed-to-expected cell number ratios (Ro/e). Values above or below 1 indicate enrichment or depletion, respectively. (**D** to **F**) Positive cell percentages (**D**), median fluorescence intensity (MFI; **E**), and IFN-γ-induced changes in MFI (ΔMFI; **F**) for surface PD-L1 and HLA-A/B/C in AGS cells and PD-L1 and MHC-I in MFC cells on soft or stiff PA hydrogels with or without IFN-γ stimulation for 24 h (n = 3). ΔMFI was calculated as stimulated MFI minus the corresponding unstimulated MFI. **P* < 0.05, ***P* < 0.01, ****P* < 0.001, *****P* < 0.0001.
